# Unleashing the Power of NR4A1 Degradation as a Novel Strategy for Cancer Immunotherapy

**DOI:** 10.1101/2023.08.09.552650

**Authors:** Lei Wang, Yufeng Xiao, Yuewan Luo, Rohan P Master, Jiao Mo, Myung-Chul Kim, Yi Liu, Urvi M Patel, Xiangming Li, Donald Shaffer, Kevin R Guertin, Emily Moser, Keiran S. Smalley, Daohong Zhou, Guangrong Zheng, Weizhou Zhang

## Abstract

An effective cancer therapy requires both killing cancer cells and targeting tumor-promoting pathways or cell populations within the tumor microenvironment (TME). We purposely search for molecules that are critical for multiple tumor-promoting cell types and identified nuclear receptor subfamily 4 group A member 1 (NR4A1) as one such molecule. NR4A1 has been shown to promote the aggressiveness of cancer cells and maintain the immune suppressive TME. Using genetic and pharmacological approaches, we establish NR4A1 as a valid therapeutic target for cancer therapy. Importantly, we have developed the first-of-its kind proteolysis-targeting chimera (PROTAC, named NR-V04) against NR4A1. NR-V04 effectively degrades NR4A1 within hours of treatment *in vitro* and sustains for at least 4 days *in vivo*, exhibiting long-lasting NR4A1-degradation in tumors and an excellent safety profile. NR-V04 leads to robust tumor inhibition and sometimes eradication of established melanoma tumors. At the mechanistic level, we have identified an unexpected novel mechanism via significant induction of tumor-infiltrating (TI) B cells as well as an inhibition of monocytic myeloid derived suppressor cells (m-MDSC), two clinically relevant immune cell populations in human melanomas. Overall, NR-V04-mediated NR4A1 degradation holds promise for enhancing anti- cancer immune responses and offers a new avenue for treating various types of cancer.

## Introduction

The tumor microenvironment (TME) consists of many cell types that cooperatively promote tumor development and progression. Most cancer therapeutics are designed to target one molecule in one defined cell type. For example, vemurafenib (BRAF inhibitor) inhibits melanoma through targeting mutated BRAF; whereas pembrolizumab (anti-PD-1 antibody) blocks the immune checkpoint PD-1 on T cells to increase anti-tumor immunity. Several FDA-approved combinations therapies involve cancer cell-killing chemotherapies and immune checkpoint inhibitors (ICI) to activate anti-cancer immunity within the TME, suggesting that co-targeting cancer cells and other cell types within the TME can be effective therapeutic regimens for cancer.

The current research focus is NR4A1, an intracellular transcription factor that is known for its crucial role not only in immune regulations but also in many other functions ^1–4^. Specifically, within the TME, NR4A1 is known to act on several cell types: 1) NR4A1 is involved in angiogenesis in the B16 melanoma model^5^. The impact of NR4A1 on neoangiogenesis has also been confirmed in several other tumor models^6–9^. As neoangiogenesis in tumors induces the formation of new blood vessels with increased permeability, NR4A1 plays a critical role in regulating basal vascular permeability by increasing endothelial nitric-oxide synthase and downregulating several junction proteins involved in adherens junctions and tight junctions^10^; 2) NR4A family has been shown to be elevated in exhausted CD8^+^ T cells, and their deletion rescues the cytotoxicity function of CD8^+^ T cell during tumorigenesis^1, 11^; 3) Tumor-infiltrating regulatory T cells (TI-Tregs) rely on NR4A1 and its other family members for their immune suppressive function^4, 12^; 4) NR4A1 can be induced in natural killer (TI-NK) cells through the IFN-γ/p-STAT1/IRF1 signaling pathway, which leads to diminished NK cell-mediated cytotoxicity against hepatocellular carcinoma^13^. Under physiological conditions, NR4A1 exerts crucial regulatory functions in B cells, limiting their expansion in response to antigen stimulation the absence of secondary signals and constraining the survival of self-reactive B cells in peripheral tissues^3^. B cells have been increasingly recognized for their important role in cancer, with studies indicating a correlation between B cell presence in the TME and improved prognosis and sensitivity to ICIs^14^. While the specific impact of NR4A1 on TI-B cells remains unexplored, the involvement of NR4A1 in modulating B cell responses underscores its potential as a therapeutic target for harnessing the anti-tumor functions of B cells in the TME.

Celastrol, a triterpenoid quinine methide derived from the root of Tripterygium wilfordii^15^, has been shown to covalently bind to NR4A1 and possesses anti-inflammatory properties^16–18^.

Celastrol is not an ideal candidate for clinical development due to limitations such as low bioavailability, a narrow therapeutic window, and undesired adverse effects^19^. Nevertheless, the moderate binding affinity of celastrol to NR4A1 (Kd of 0.29 μM)^20^ makes it a promising starting point as a warhead for the design and synthesis of NR4A1 proteolysis-targeting chimeras (PROTACs). PROTACs are bifunctional molecules that consist of a pharmacophore targeting a protein of interest and another pharmacophore recruiting an E3 ligase such as von Hippel Lindau (VHL) or cereblon (CRBN), facilitating polyubiquitination and subsequent proteasome degradation of the target protein^21, 22^. PROTAC represents a novel approach in drug discovery. Unlike the conventional “occupancy-driven” inhibitor-based therapy, PROTACs work in a catalytic and “event-driven” manner, leading to the depletion of protein targets. This unique mechanism of action offers several advantages including higher potency, prolonged pharmacological effect, and improved selectivity. More importantly, PROTACs offer new possibilities to target traditionally undruggable or difficult-to-drug targets, such as transcriptional factors, scaffolding proteins, and multi-functional proteins that cannot be effectively inhibited by a single inhibitor.

Here, we report a first-of-its-kind NR4A1-targeting PROTAC named NR-V04 as a lead for cancer therapy. We have performed extensive characterization of NR-V04, which reveals its promising pharmacokinetic and pharmacodynamic profile, degradation efficiency and specificity for NR4A1, tumor-inhibitory effect and immune activation within the TME, along with its low toxicity within the therapeutic dosing range.

## Results

### NR4A1 is elevated in many cell types within the TME of human cancers and is a valid target in the TME for cancer therapy

We recently found that *NR4A1* is the most elevated gene in the TI-Tregs from human renal clear cell carcinoma (RCC, GEO: GSE121638)^23^. We further analyzed *NR4A1* expression within several single-cell RNA sequencing datasets (scRNAseq), including human hepatocellular carcinomas (HCC, GEO: GSE98638)^24^ and melanomas (GEO: GSE120575)^25^. In HCC, we confirmed that *NR4A1* expression was significantly elevated in TI-Tregs relative to Tregs from blood or normal tissues, conventional CD4^+^ T cells (Tconv), or CD8^+^ T cells (Fig. 1A, note the difference in FPKM scales). Exhausted Tconv or CD8^+^ T cells also had increased levels of *NR4A1*, though at much lower levels than that in TI-Tregs (Fig. 1A). In melanoma, there is diverse expression pattern of *NR4A1* across multiple cell types within the TME, but most TI immune cell types showed elevated *NR4A1* expression than their blood counterparts (Fig. 1B). *NR4A1* expression was particularly evident in B cells, monocytes or macrophages (M), dendritic cells (DCs), Treg cells, and exhausted CD8^+^ T (Texh) cells (Fig. 1C)^25^. *NR4A2* displayed a ubiquitous presence across all immune cell populations, while *NR4A3* predominantly appeared in monocytes or macrophages, DCs, and a specific subset of CD8 T cells (Fig. 1C). Leveraging the TCGA datasets, we investigated NR4A1 expression in melanoma patients and identified a negative correlation between NR4A1 and anti-tumor immune responses, including the gene expression of *IFN*γ (interferon γ), *GZMB* (granzyme B), and *PRF1* (perforin) (Supplementary Fig. 1A-1C). Gene set enrichment analysis (GSEA) revealed that *NR4A1* expression was inversely associated with T cell receptor (TCR) and B cell receptor (BCR) signaling pathways (Supplementary Fig. 1D-1G). Collectively, these findings highlight the potential involvement of NR4A1 in the immune modulation of human cancers.

**Figure 1.**
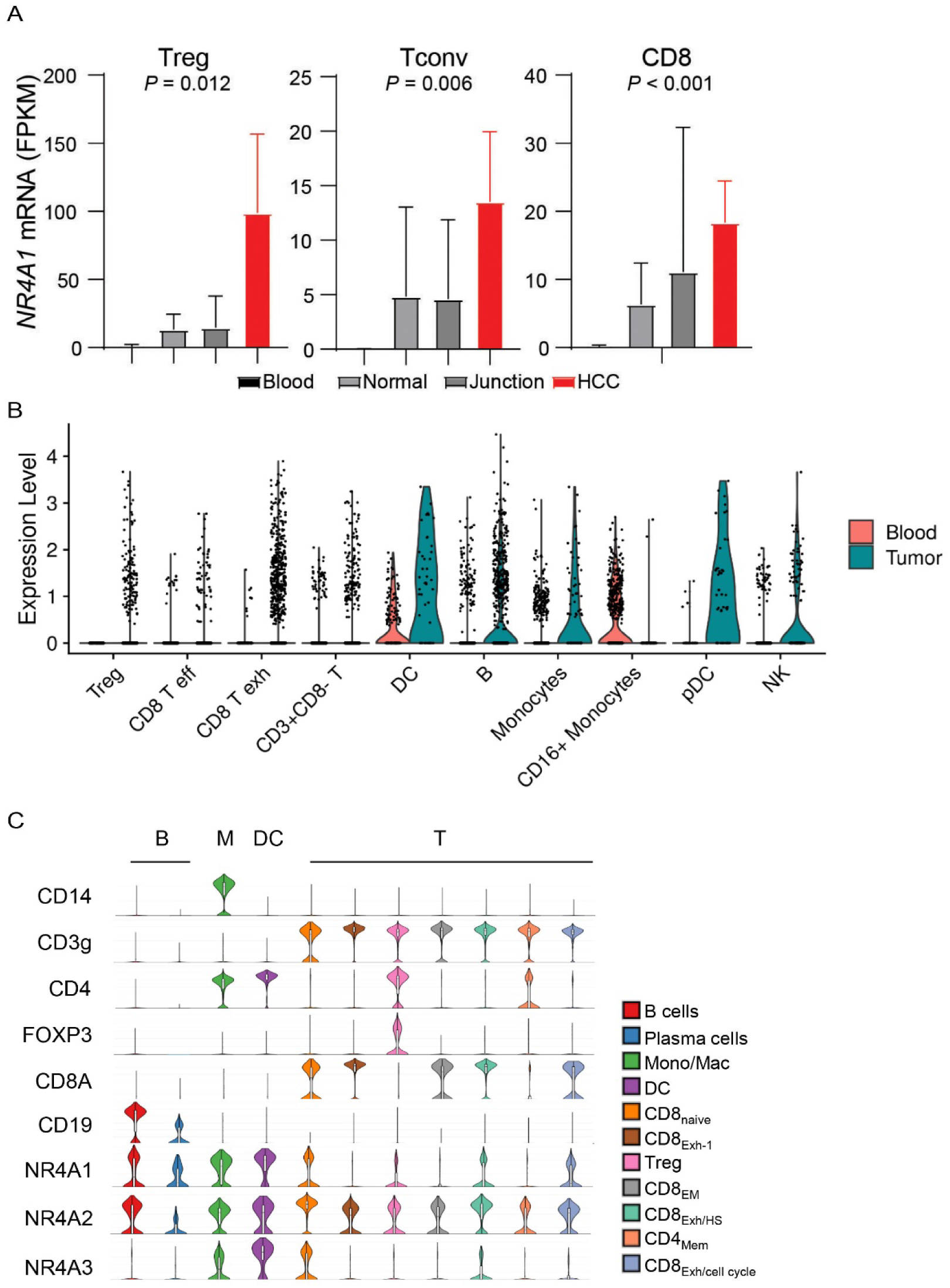
*NR4A1* is elevated in several tumor-promoting immune cells within the tumor microenvironment. **A.** Expression of *NR4A1* in T cells from blood, normal parenchyma, adjacent normal junction, or human hepatocellular carcinomas (HCC, GSE98638). **B.** Violin plots showing *NR4A1* expression in different immune cells from human blood or melanomas. **C.** Violin plots showing gene expression including NR4A family and lineage markers in human melanomas (GSE120575).

To determine whether NR4A1 plays important roles in the TME, we implanted 3 syngeneic tumors into wild-type (WT) or *NR4A1*^-/-^ (KO) mice, including MC38 colon cancer model, Yummer1.7 and B16F10 melanoma models. We used a minimal number of tumor cells that can produce tumors in WT mice and found that *NR4A1*^-/-^ (KO) mice exhibited much slower tumor growth rate in all of the tumor models (Fig. 2A-2C). MC38 model showed very minimal tumor growth that peaked at the 3^rd^ week in the KO mice but was regressed thereafter (Fig. 2A), suggesting that tumors be eradicated by the induction of anti-tumor immunity in KO mice. These data strongly support the role of NR4A1 in the TME and immune modulation, and suggest that targeting NR4A1 is a potentially promising immunotherapy for cancer.

**Figure 2.**
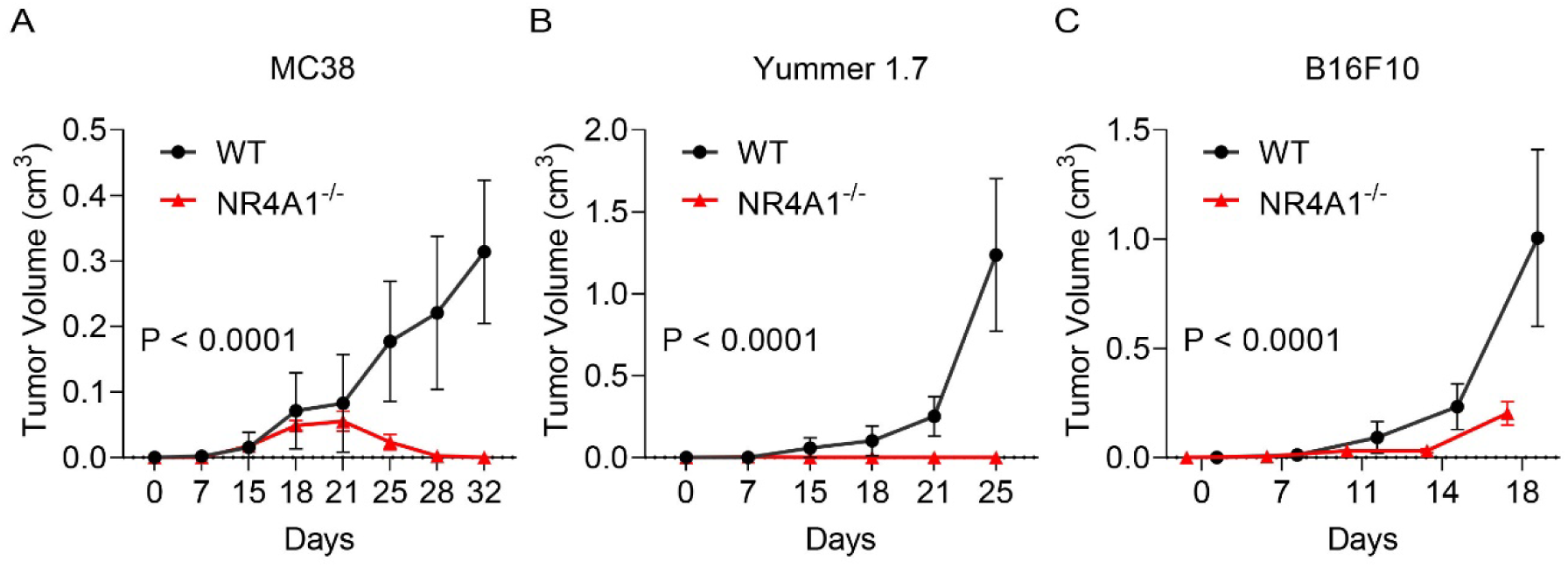
NR4A1-deletion leads to diminished tumor growth. **A-C.** Primary tumor growth curve in littermates of wild type (WT) or *NR4A1*^-/-^ mice, including (**A**) MC38 colon cancer, (**B**) Yummer1.7 melanoma and (**C**)B16F10 melanoma. n = 7-8 mice per group.

### The design and screening of PROTACs against NR4A1

A number of NR4A1 ligands have been reported^26^, and celastrol is one of the few that has been well characterized. Celastrol covalently binds to NR4A1 through engaging cysteine C551, and the binding affinity is within the sub-nanomolar range based on several biophysical assays^16–18^. The structure and activity relationship (SAR) study of celastrol on NR4A1 has been explored, which indicates that the carboxylic acid group of celastrol is amenable for chemical modifications ^17^. Therefore, we reasoned that celastrol might be a suitable warhead for PROTAC construction. We performed a molecular docking study between celastrol and the ligand binding domain (LBD) of NR4A1 and found that the carboxylic acid group is solvent-exposed, which represents an ideal tethering site for linker attachment (Fig. 3A). We utilized polyethylene glycol (PEG) linkers for the conjugation of celastrol to a VHL E3 ligase ligand via two amide bonds. As a proof-of-concept study, three PROTACs with different PEG linker lengths were synthesized (Fig. 3B) and tested in CHL-1 human melanoma cells (Fig. 3C-3D). Celastrol treatment did not alter the protein level of NR4A1 (Fig. 3C-3D). NR-V04 – which bears a 4-PEG linker, exhibited the highest potency in reducing the protein level of NR4A1 (Fig. 3C-3D) and was thus chosen for further investigations.

**Figure 3.**
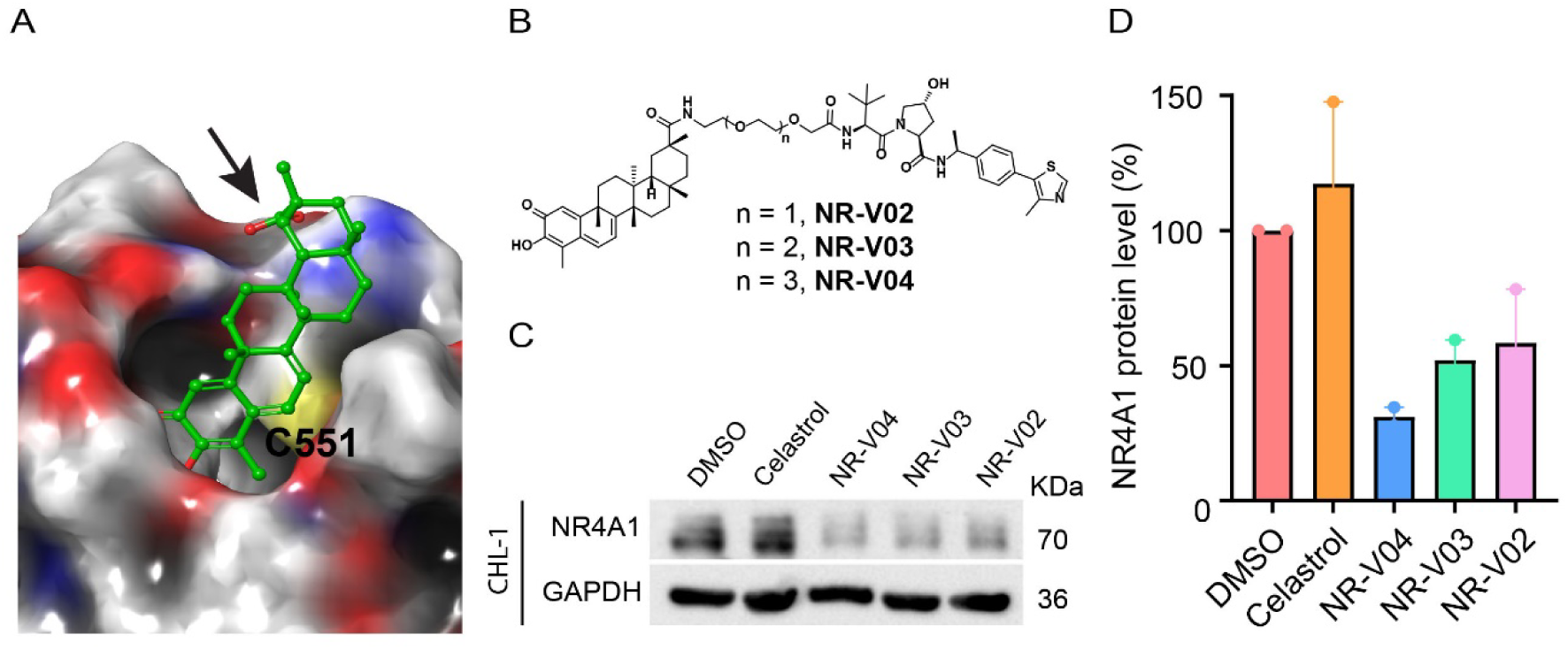
The design and synthesis of NR4A1 PROTACs. **A**. Docking study revealed the carboxylic acid in celastrol is a potential tethering vector for PROTAC construction. **B**. The structure of synthesized PROTACs. **C-D**. The initial screening of NR4A1 degradation in CHL-1, a human melanoma cell line. CHL-1 cells were treated with 250 nM PROTACs for 16 hours and the degradation was determined by **(C)** immunoblotting and **(D)** densitometry. n=2.

### NR-V04 efficiently reduces NR4A1 protein level

Following a 16-hr treatment, NR-V04 induced a dose-dependent decrease of NR4A1 protein in CHL-1 cells with a 50% degradation concentration (DC_50_) of 228.5 nM and A375 melanoma cells with a DC_50_ of 518.8 nM (Fig. 4A). NR-V04 treatment also reduced NR4A1 protein level in WM164 and M229 human melanoma cells (Supplementary Fig. 2A), and SM1 and SW1 – two mouse melanoma cell lines (Supplementary Fig. 2B). Celastrol did not change NR4A1 protein level (Fig. 4B). Notably, NR- V04 treatment led to accumulation of the VHL protein (Fig. 4A and Supplementary Fig. 2A).

**Figure 4.**
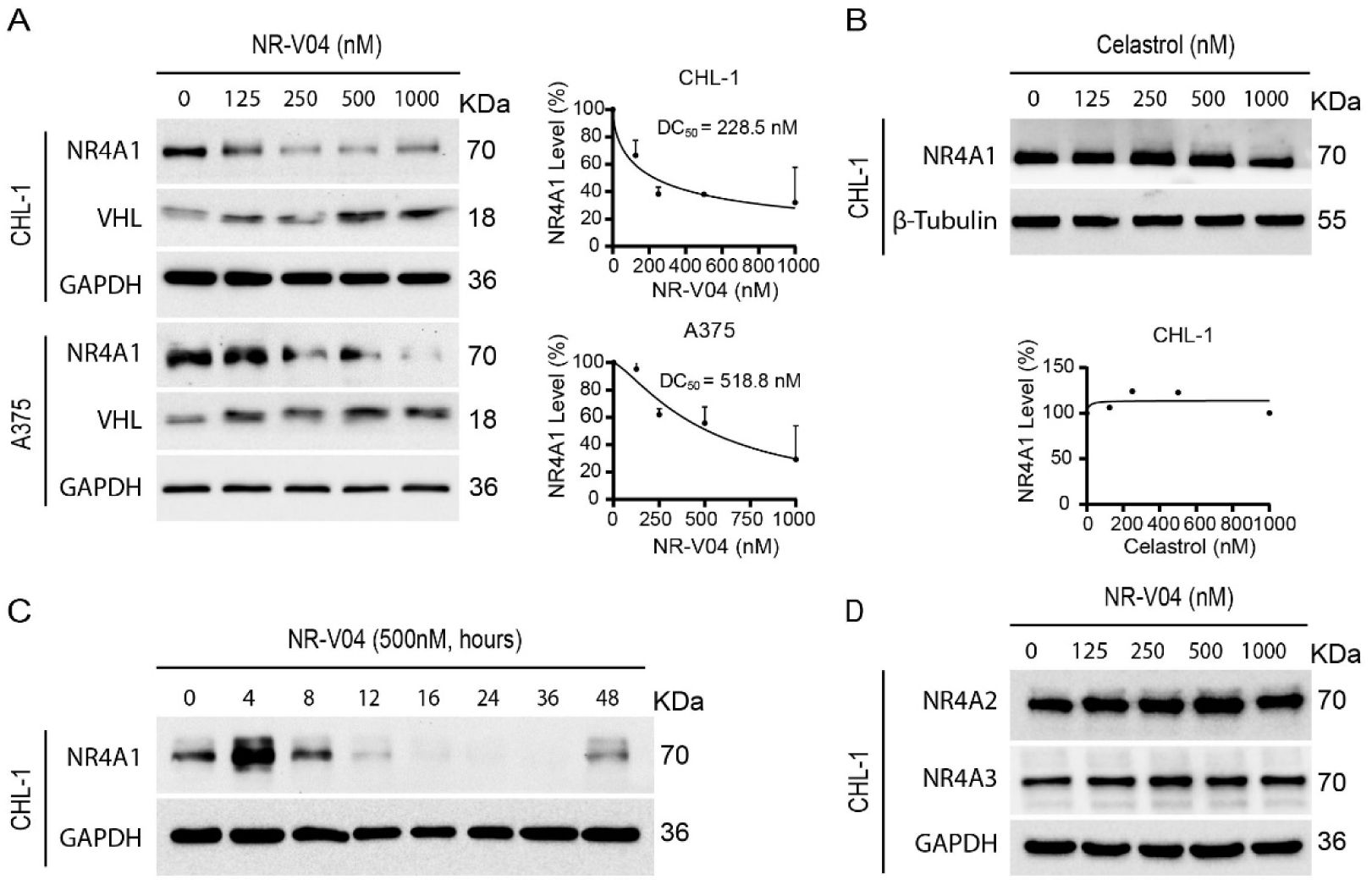
NR-V04 induces NR4A1 degradation. **A**. NR-V04 effectively promoted the degradation of NR4A1 in two human melanoma cell lines in 16 hours, CHL-1 (DC50 = 228.5 nM) and A375 (DC50 = 518.8 nM), while simultaneously stabilizing VHL protein. **B**. Celastrol treatment did not result in any significant change in the expression level of NR4A1 in the CHL-1 cell line in 16 hours. **C**. Time-dependent degradation of NR4A1. CHL-1 cells were treated with 500nM of NR-V04 at the indicated time points. **D.** NR-V04 did not induce the degradation of NR4A2 and NR4A3. A-D: n=2.

Time course studies indicated that the efficient reduction of NR4A1 occurred between 8-48 hrs after NR-V04 treatment (Fig. 4C). Interestingly, we observed an initial induction of NR4A1 protein after 4 hrs of NR-V04 treatment (Fig. 4C), along the timeline of increased *NR4A1* mRNA after 4 hrs of NR-V04 treatment (Supplementary Fig. 3). We reasoned that this induction should be mainly caused by celastrol, because we observed an increase in NR4A1 mRNA levels after 2 hrs of celastrol treatment (Supplementary Fig. 3). Among the three NR4A family members, NR-V04 selectively reduced the NR4A1 protein level while sparing NR4A2 and NR4A3 (Fig. 4D, Supplementary Fig. 2C). Our data support that NR-V04 is an effective NR4A1 degrader *in vitro*.

### NR-V04 induces ternary complex formation and proteasome-mediated NR4A1 degradation

Ternary complex formation is a prerequisite for PROTAC to mediate protein degradation ^22^. We employed proximity ligation assays (PLA) to detect localized signals only when NR4A1 and VHL are in close proximity with the presence of NR-V04. NR-V04 treatment induced very strong PLA signals but not when cells were treated with celastrol or DMSO (Fig. 5A, Supplementary Fig. 4A). Additionally, we conducted co-immunoprecipitation (Co-IP) experiment using Flag-NR4A1 expressed in the HEK293T cells with and without NR-V04 treatment. Notably, NR-V04 treatment led to the formation of a complex between NR4A1 and VHL, which was not observed with DMSO-treated cells (Fig. 5B).

**Figure 5.**
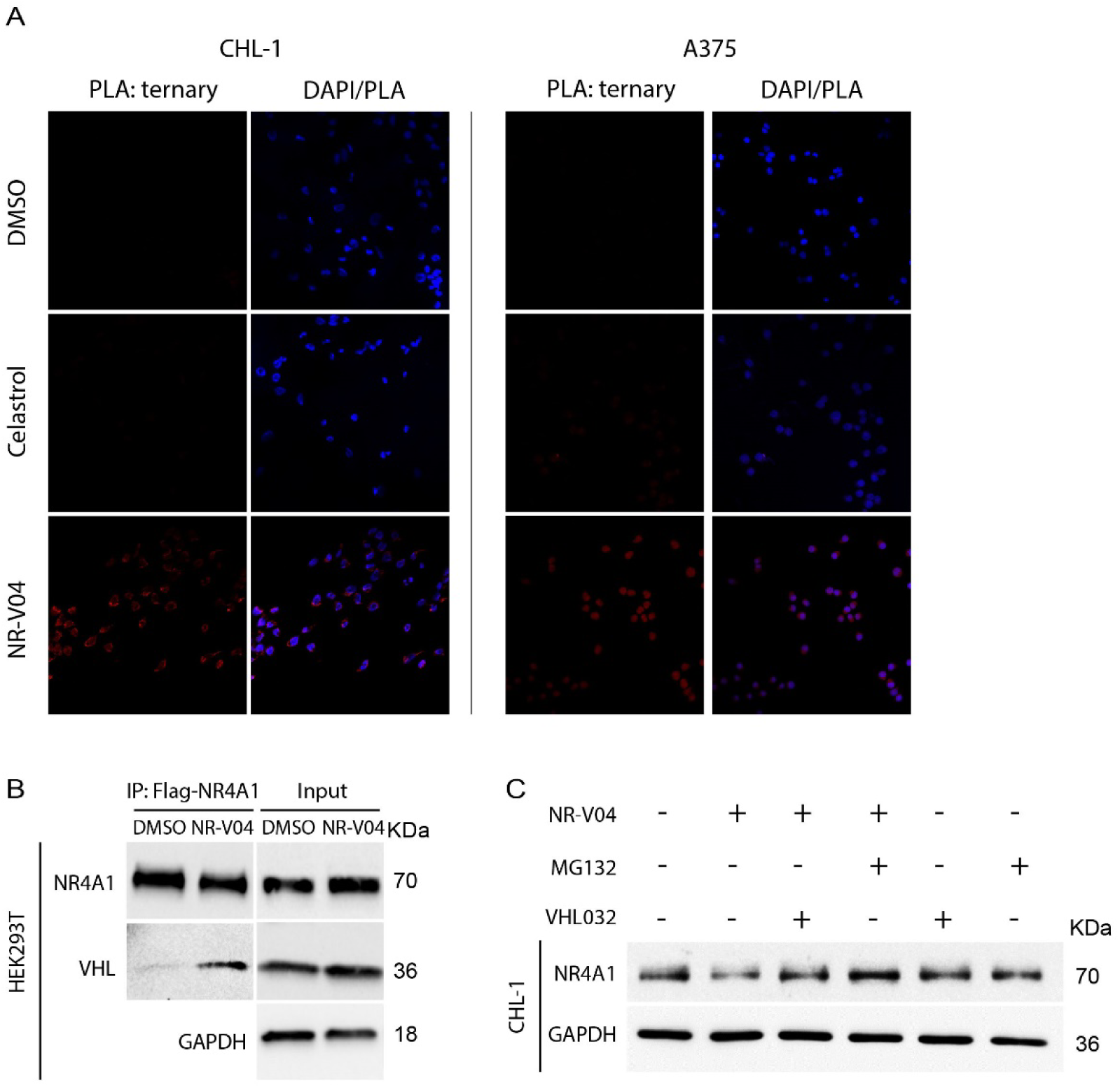
NR-V04 induces a ternary complex formation and mediates NR4A1 degradation through the ubiquitin-proteasome system. **A**. Proximity Ligation Assay (PLA) showing ternary complex formation induced by NR-V04. CHL-1 (left panels) and A375 (right panels) cells were treated with DMSO, 500 nM Celastrol, or 500 nM NR-V04 for 16 hrs. Representative images were shown for PLA assay (20 x magnification). **B**. Co-immunoprecipitation (co-IP) experiment showing complex formation between NR4A1 and VHL by NR-V04 treatment. Co-IP was performed in NR4A1-Flag overexpressed HEK293T cells that were pretreated with 0.5 μM MG132 for 10 minutes, followed by 16-hour treatment with DMSO or 500 nM NR-V04. NR4A1 was pulled down using an anti-Flag antibody conjugated to magnetic beads. **C**. NR-V04 induces NR4A1 degradation via VHL E3 ligase- and proteasome-dependent manner. CHL-1 cells were pretreated with 0.5 μM MG132 or 10 µM VHL 032 for 10 minutes, followed by 16-hour treatment with DMSO or 500 nM NR-V04. A-C: n=2.

We further determined the mechanism of action (MOA) of NR-V04-mediated reduction of NR4A1 protein. MG132, a proteasome inhibitor, effectively rescued NR4A1 protein level reduced by NR-V04 (Fig. 5C), which supports that NR-V04-induced reduction of NR4A1 protein is through the proteasome pathway. VHL-032, the VHL ligand used for NR-V04 construction, prevented the NR-V04-induced degradation of NR4A1 (Fig. 5C), which is further supported using the VHL-knockout cells (Supplementary Fig. 4B).

### NR-V04 has good pharmacokinetic (PK) property and shows prolonged NR4A1 degradation in vivo

Our mouse study showed that NR-V04 had a half-life of 8.6 hrs via intraperitoneal injections (*i.p.*) injection and 5.36 hrs via the *i.v.* injection and near complete absorption (F=98%) through *i.p.* administration with a low clearance rate (Fig. 6A). Encouraged by the PK result, we assessed the *in vivo* pharmacodynamic properties (PD) of NR-V04. Mice bearing MC38 tumors were administered two effective doses of NR-V04 (1.8 mg/kg as determined by a pilot dose-dependent experiment, data not shown), and the tumor tissues were collected and analyzed 1 to 4 days after the last treatment. NR-V04 treatment led to significant degradation of NR4A1 starting from day 1, and persisted for 3 days, with a slight recovery of NR4A1 on day 4 (Fig. 6B). Based on the PD result, the dose regimen was set at 1.8 mg/kg, twice a week for following tumor studies. We also analyzed NR4A1 protein in MC38 tumors with mice treated with vehicle, celastrol and NR-V04 after a total of 8 doses for 4 weeks in tumor-bearing mice. The data showed that NR4A1 was almost completely degraded in the NR-V04-treated group, but not in the celastrol- or vehicle-treated group (Fig. 6C).

**Figure 6.**
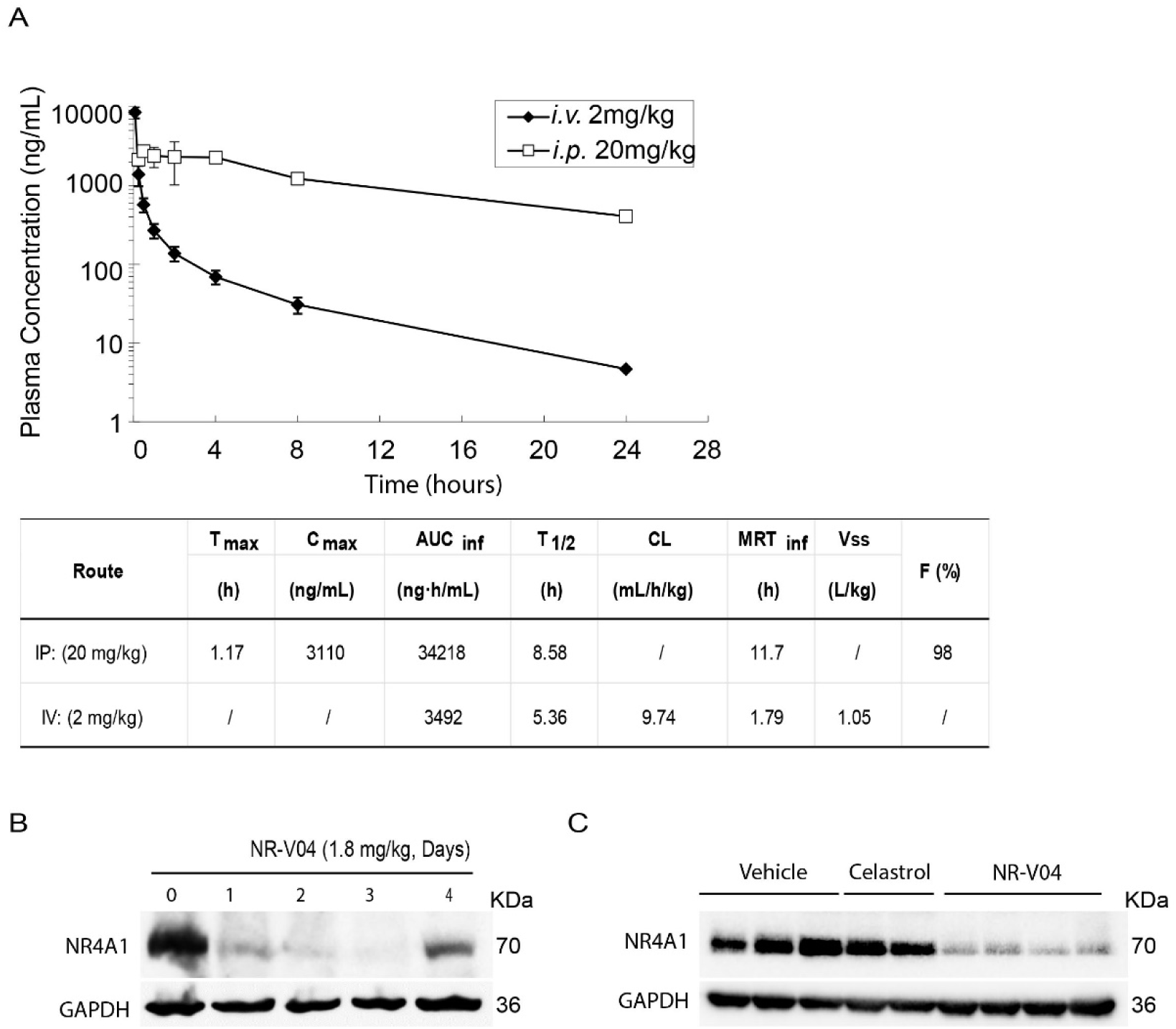
The pharmacokinetic (PK) and pharmacodynamic (PD) study of NR-V04. **A**. PK parameters of NR-V04 in WT mice. n = 3. **B**. NR-V04 induces long-lasting degradation of NR4A1 in MC38 tumors. Mice bearing MC38 tumors were treated with two-dose administration of NR-V04 via *i.p.* injection at 1.8 mg/kg, and tumors were collected at indicated timepoints. Tumor lysates were analyzed by immunoblotting with each lane representing an individual tumor lysate. n=2 **C**. Immunoblotting showing NR4A1 degradation in MC38 tumors upon termination. MC38 tumor-bearing mice were treated with vehicle, 1.8 mg/kg NR-V04 or 0.75 mg/kg celastrol treatment (equivalent to 1.67 μmol/kg), every 4 days until experimental end points. Tumor tissues were collected for lysate collection and immunoblotting (n = 4).

### NR-V04 inhibits tumor growth

Based on the PK and PD data, we conducted iterative optimization of NR-V04 treatment in tumor-bearing mice. Our final treatment regimen was set at a dose of 1.8 mg/kg NR-V04 or 0.75 mg/kg celastrol (equivalent to 1.67 μmol/kg) via the *i.p.* routes, administered twice a week. Treatment was initiated 7 days after inoculation of cancer cells when tumors were palpable. NR-V04 treatment effectively inhibited tumor growth in MC38 model (Fig. 7A) and Yummer1.7 melanoma generated from a tumor formed in a *Braf*^V600^/*Cdkn2a*^-/-^/*Pten*^-/-^ mouse (Fig. 7B), as well as the most commonly used B16F10 melanoma model (Fig. 7C). In contrast, celastrol treatment resulted in either increased growth of B16F10 tumors (Supplementary Fig. 5A) or failed to inhibit tumor growth of MC38 tumors (Supplementary Fig. 5B). To validate NR4A1 as the primary molecular target of NR-V04 in the TME, we inoculated B16F10 melanoma tumors in *NR4A1*^-/-^ mice and found that NR-V04 failed to inhibit B16F10 in these mice (Fig. 7D). *Nod*/*Scid*/*Il2rg*^-/-^ (NSG) mice lack T cells, B cells, and NK cells and NR-V04 treatment failed to inhibit tumor growth in the NSG hosts for MC38 or B16F10 tumors (Fig. 7E-7F).

**Figure 7.**
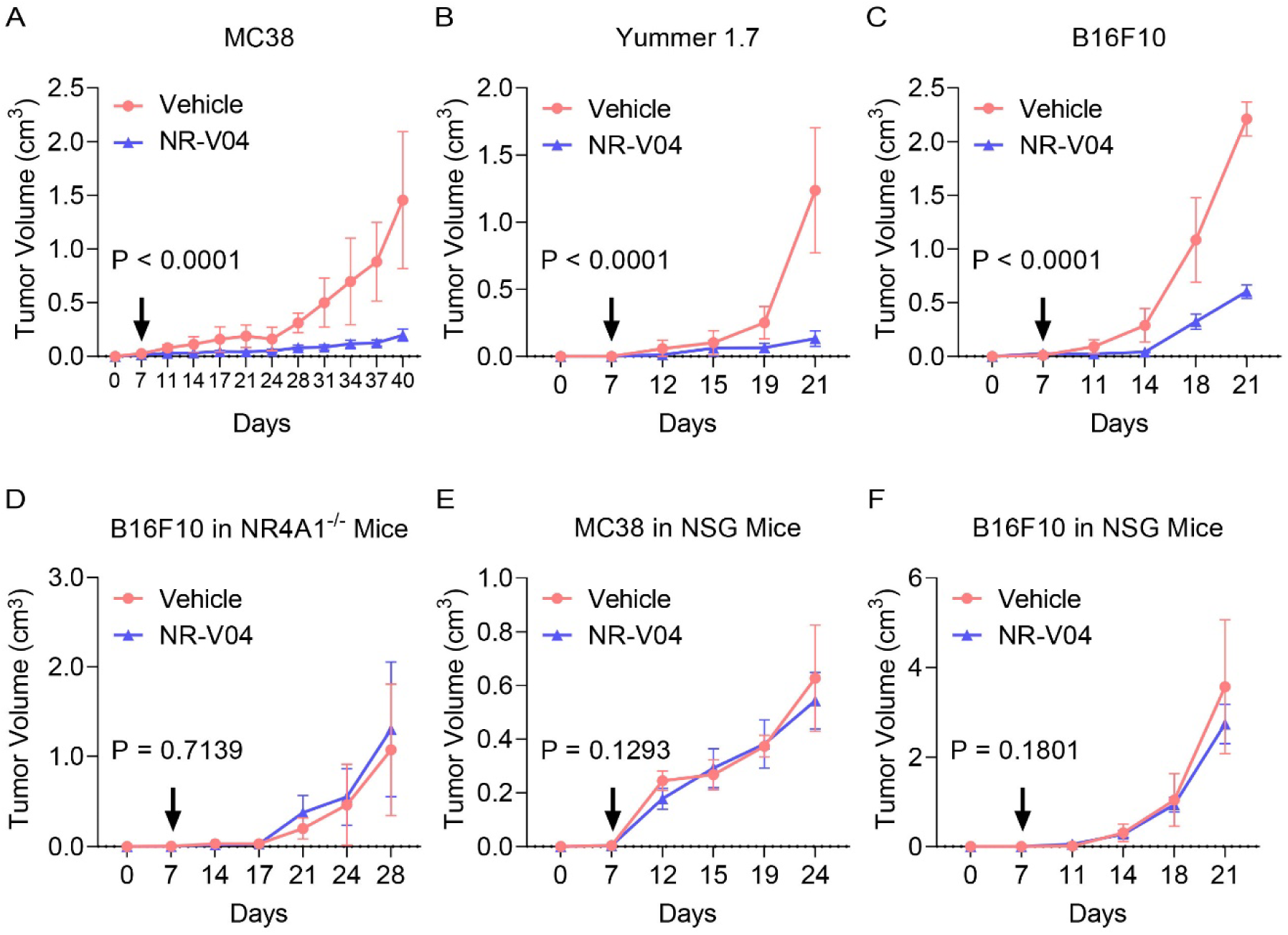
NR-V04 exhibits outstanding anti-tumor effects in several tumor models. **A-F**. Tumor-bearing mice were treated with 1.8 mg/kg NR-V04 or vehicle via *i.p.* injection on day 7 when tumors were palpable. The treatment was repeated every four days until the tumors reached the endpoint size of 2 cm in diameter. **(A**) NR-V04 inhibited MC38 colon adenocarcinoma growth, n=4 per group; **(B**) NR-V04 inhibited Yummer 1.7 melanoma growth, n=4 per group. (**C**) NR-V04 inhibited B16F10 melanoma growth, n=5 per group. **(D**) NR-V04 failed to inhibit B16F10 melanoma growth in *NR4A1*^-/-^ mice, n=5 per group. (**E**) NR-V04 failed to inhibit MC38 colon adenocarcinoma growth in NSG mice, n=4. (**F**) NR-V04 failed to inhibit B16F10 melanoma in NSG mice, n=5. Two-way ANOVA was performed for all tumor growth curves with *P* values indicated.

### NR-V04 modulates different immune responses within the TME

As we were aiming to understand the immune regulatory effects of NR-V04, we designed studies to minimize the influence of tumor size on the tumor immune microenvironment. We allowed tumors to grow to 1 cm in diameter and treated tumor-bearing mice with 2 doses of NR-V04 at similar dose/time intervals (on day 1 and day 4). Tumors were collected on day 7. In the B16F10 model, NR-V04 significantly increased TI-B cells (Fig. 8A showing B220^+^ cells from 14.7% to 30.1% and gating scheme shown in Supplementary Fig. 6), whereas no significant change was observed in the spleen and blood. Furthermore, we identified the increased portion as CD38^+^CD138^-^ plasmablasts, and the BCR isotypes are either IgD^+^IgM^-^ or IgD^+^IgM^+^ (Supplementary Fig. 7A-7B).

**Figure 8.**
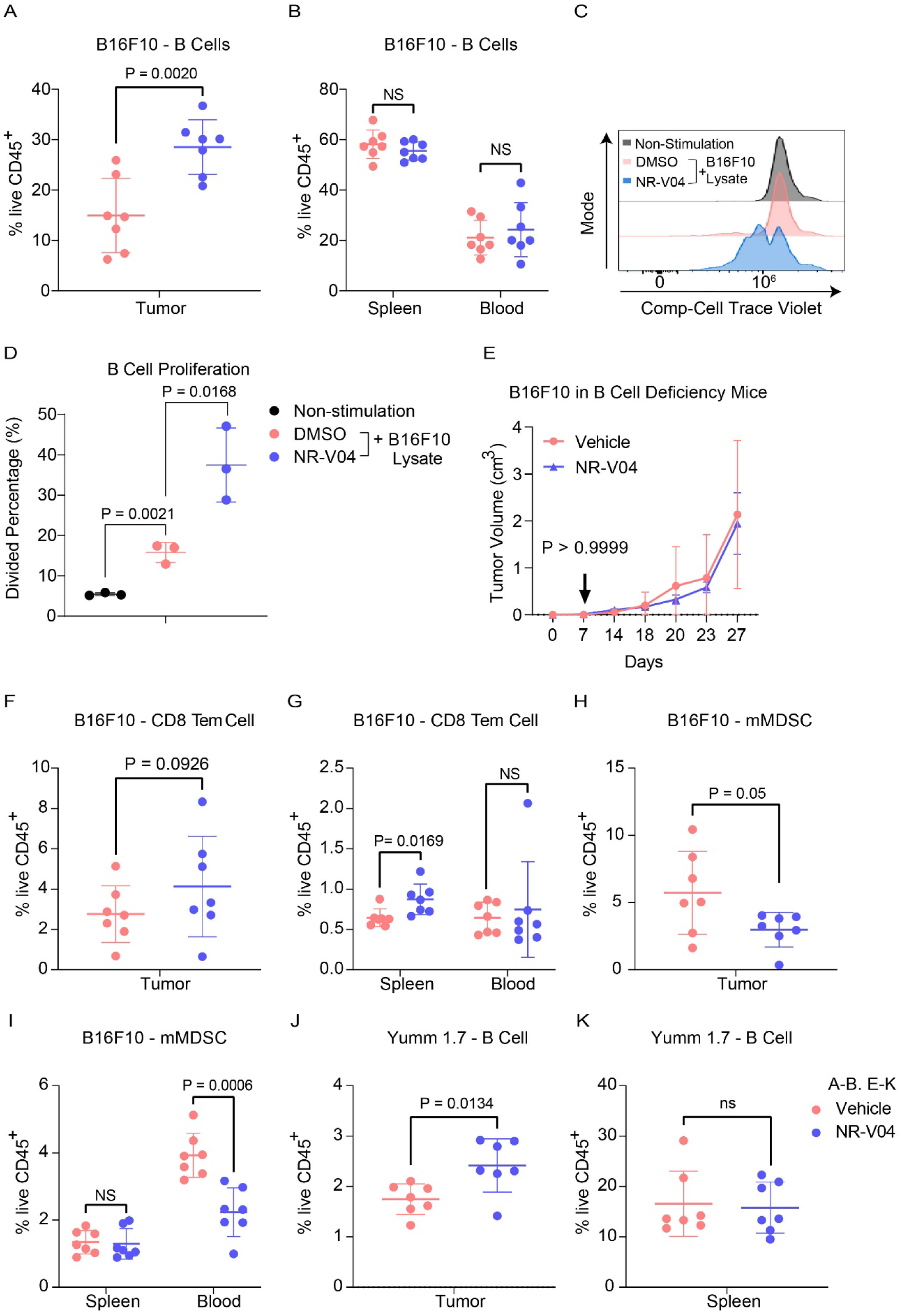
Effect of NR-V04 on immune cells in the TME. **A-B and E-K**. Tumor-bearing mice were treated with 1.8 mg/kg NR-V04 or vehicle via *i.p.* injection when tumor size reached 1 cm in diameter, with two treatments started as day 1 and repeated on day 4. Tumors were collected and single cells were isolated from tissues for flow cytometry analysis. **A-B**. NR-V04 treatment increases B cell percentage in the TME, but not in spleen and blood in mice inoculated with B16F10 melanoma, n=7. **C-D**. NR4A1 depletion induces B cell proliferation. B cells isolated from spleen were labeled with cell trace violet, untreated or treated with B16F10 lysis, following with the cotreatment of DMSO or 500 nM NR-V04 for 24 hours (n=3). B cell proliferation was determined by flow cytometry. **E**. NR-V04 fails to inhibit B16F10 melanoma growth in mice deficient of mature B cells (n=5). **F-G**. NR-V04 treatment increased effector memory CD8 T cell percentage in spleen, but not in tumors and blood in B16F10 melanoma (n=7). **H-I**. NR-V04 treatment decreased mMDSC percentage in tumor and blood, but not in spleen in B16F10 melanoma, n=7. **J-K**. NR-V04 treatment increased B cell percentage in the TME, but not in spleen and blood in Yumm 1.7 melanoma (n=7). **E**. Two-way ANOVA was performed for the tumor growth curve with *P* values indicated. Others are shown as the mean ± SD. A two-sided unpaired *t* test was performed, with *P* values indicated.

B cells from the spleen of *NR4A1*^-/-^ mice exhibited increased proliferation (72.8%) compared to those from WT mice (35.2%) upon 10 μg/mL anti-IgM stimulation (Supplementary Fig. 8A-8B), in agreement with published literature showing that NR4A1 is involved in limiting B cell proliferation^3^. Since there is no report related to NR4A1 in TI-B cells, we treated WT B cells with B16F10 lysates to induce B cell proliferation, with or without NR-V04 treatment. NR-V04 led to NR4A1 degradation in B cells (Supplementary Fig. 8C-8D) and significantly increased B cell proliferation (Fig. 8C-8D). We inoculated B16F10 into B6.129S2-*Ighm*^tm1Cgn^/J strain mice that lack mature B cells and found that NR-V04 was unable to inhibit tumor growth (Fig. 8E), supporting that the tumor inhibitory function of NR-V04 is mediated through B cell elevation in the TME.

We also observed significant increases in memory effector CD8 T cells (CD44^+^CD62L^-^ Tem, Fig. 8F-8G) in the spleen and a decrease in myeloid-derived monocytic suppressor cells (mMDSC, CD11B^+^Ly6C^+^, Fig. 8H-8I) in the blood upon NR-V04 treatment.

We investigated the immune profile in the Yumm1.7 tumor model, known for its human-relevant genetic mutations in melanoma cancer, including *Braf*^V600E/wt^/*Pten*^-/-^/*Cdkn2*^-/-27^. NR-V04 treatment also resulted in a significant increase in the TI-B cells (Fig. 8J-K), supporting a general mechanism of action for NR-V04 in TI-B cell proliferation and induction.

### NR-V04 exhibits an excellent safety profile in mice

Using tumor-bearing mice (Fig. 9), NR-V04 did not significantly induce body weight changes, nor did it induce significant changes in peripheral blood or spleen (Fig. 9). We further assessed the toxicity of NR-V04 using both male and female C57BL/6J mice with increased doses up to 5 mg/kg (Fig. 9A); there were no significant change in body weight (Fig. 9B). Complete blood counts (CBC) were determined at different time points afterNR-V04 treatment that did not result in any significant changes of the hematology parameters, including whole blood cell count (Fig. 9C), lymphocytes (Fig. 9D), neutrophils (Fig. 9E), red blood cells (Fig. 9F), platelets (Fig. 9G), mean platelet volume, mean corpuscular volume, hemoglobin concentration, mean corpuscular hemoglobin, hematocrit percentage, and mean corpuscular hemoglobin concentration (Supplementary Fig. 9A-9F).

**Figure 9.**
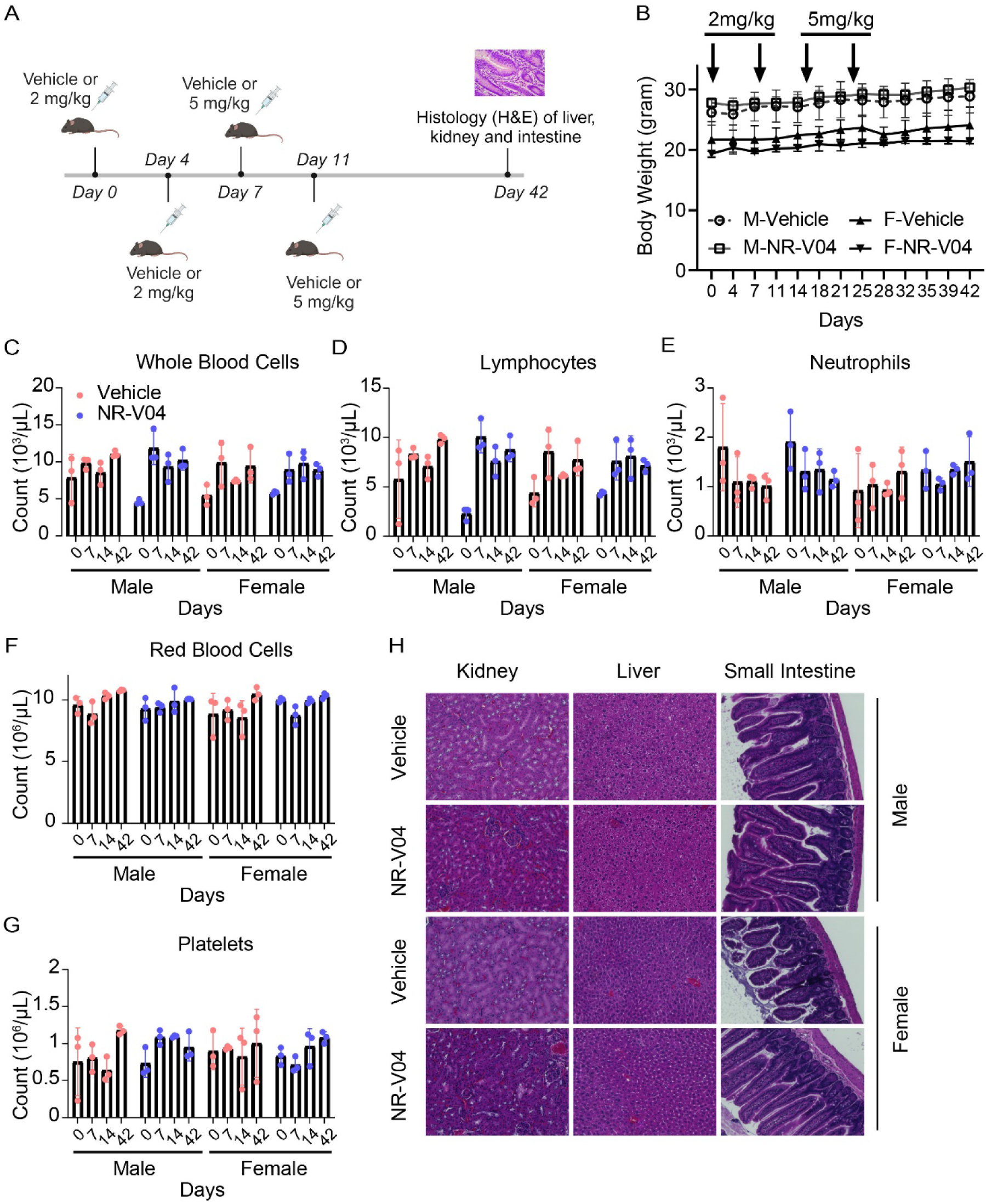
NR-V04 has minimal toxicity. **A**. Schematic of the toxicity testing. Male and female mice were treated with two doses of 2 mg/kg NR-V04 and two doses of 5 mg/kg NR-V04 over two weeks. Blood samples were collected on day 0, 7, 14, and 42 for hematology analysis, and body weight was measured twice per week. After day 42, all mice were euthanized, and tissues (kidney, liver, and small intestine) were harvested for H&E staining (n=3). **B**. Mice did not experience significant weight loss with NR-V04 treatment during the 42-day period. **C-G**. Hematology analysis of different blood cell components after NR-V04 or vehicle treatment, including **(C)** whole blood cells, **(D)** lymphocytes, **(E)** neutrophils, **(F)** red blood cells, and **(G)** platelets. **H.** NR-V04 impacts on tissue histology, including representative kidney, liver, and small intestine.

Furthermore, we conducted a histological examination using hematoxylin and eosin (H&E) staining to evaluate the kidney, liver, and small intestine tissues in both male and female mice at day 42. Notably, NR-V04 treatment did not induce tissue damage in any of these organs (Fig. 9H). These findings provide important insights into the potential clinical application of NR-V04 as a safe and effective immunotherapeutic agent.

## Discussion

In this study, we successfully developed NR-V04, a PROTAC that efficiently degrades NR4A1 in the TME. NR-V04 has strong anti-tumor effects via targeting various cell types, including T cells, B cells, and MDSCs. Though the cancer cell-intrinsic roles of NR4A1 have been extensively characterized in previous studies^28–30^, our current study strongly supports that NR4A1 plays critical roles in immune modulation within the TME. As such, we have established NR4A1 as a valid target for cancer immunotherapy.

We have discussed several potential cellular targets by NR4A1 degradation, including T cells, NK cells, and endothelial cells etc., but we hadn’t expected TI-B cells to be the most significantly induced cell type by NR-V04 treatment. The role of B cells within the TME is complex and has been debated depending on cancer types, B cell subpopulations, and/or the contrasting functions of TI-B cells. On the one hand, higher B cell numbers – often associated with the tertiary lymphoid structures (TLS), are associated with favorable clinical outcomes and improved responsiveness to immunotherapies in melanoma ^31, 32^; on the other hand, regulatory B (Breg) cells can mediate immunosuppression, and certain B cell subpopulations can promote tumor growth through pro-inflammatory mechanisms ^33, 34^. In our study, NR-V04 significantly increased B cell numbers in B16F10 and Yumm 1.7 mouse melanoma models, and B cell deficiency completely abolishes the tumor inhibitory effect of NR-V04. The importance of B cells in dampening B16F10 melanoma growth has been supported by previous research^35^, further underscoring the significance of NR-V04-mediated B cell activation. We investigated the mechanistic underpinnings of NR-V04’s impact on B cells and found that NR4A1 expression limits B cell responsiveness to tumor antigens, in agreement with a previous study^3^. B cells play two critical roles during tumor development: 1) secreting antibodies targeting specific tumor antigens^36–40^ and 2) presenting tumor antigens to activate the T cell response^41–44^. With NR-V04 treatment, we observed an increase in the B cell population that primarily consists of plasmablasts, a subset of B cells known for their rapid production and early antibody responses to tumor antigens^45^. This increase in plasmablasts is associated with a favorable prognosis and has been observed within tumors^31^. Additionally, NR-V04 treatment enhances the expression of both BCR isotypes, IgD^+^IgM^-^ and IgD^+^IgM^+^, suggesting an enhanced B cell response to tumor antigens. An additional advantage of NR-V04’s B cell regulation is its selective impact on the TME, sparing the B cells or other immune cells in the spleen and blood. This minimizes potential side effects, such as autoimmune responses triggered by B cell antibodies in peripheral tissues. It is crucial to acknowledge that the frequency of B cell infiltration varies among different tumor types. For example, B16F10 melanoma exhibits a high B cell infiltration, accounting for more than 50% of total tumor-infiltrating lymphocytes, whereas Yumm 1.7 shows a much lower B cell infiltration with only 2-3% of total tumor-infiltrating lymphocytes. This variation directly correlates with NR-V04’s therapeutic outcomes, implying that NR-V04 application may achieve more favorable responses in cancers with high B cell infiltrations.

NR-V04’s impact on the TME extends beyond regulating B cells to influence anti-tumor immunity. NR-V04 decreases mMDSCs (CD11B^+^Ly6C^+^) in tumors and blood. mMDSCs are known to suppress the immune response, including B cell function^46, 47^. Furthermore, B cell activation results in immune complex formation that attracts pro-inflammatory cytokines produced by mMDSCs^31, 48^. Thus, NR-V04’s action on mMDSC can potentially alleviate immune suppression, thereby enhancing B cell-mediated anti-tumor effects. Moreover, NR-V04 significantly increases CD8^+^ Tem cells in the spleen of B16F10 melanoma-bearing mice, supporting potential systematic protection of tumor cell recurrence when encountered with those CD8+ Tem cells.

The PK-PD decoupled, long-lasting degradation effect of NR-V04 *in vivo* (Fig. 6B) is expected for a PROTAC molecule but could also suggest that NR-V04 can specifically accumulate in the tumors, a favorable feature for drug development that warrants reasonable lower effective doses and longer treatment intervals. The warhead used in NR-V04 is celastrol which has been associated with several adverse effects such as hepatotoxicity^49, 50^, hematopoietic system toxicity^51^, nephrotoxicity^52^, weight loss, and negative impact on metabolic and cardiovascular functions^53–55^. However, with the same treatment regimen, NR-V04 exhibits an excellent safety profile *in vivo*. This safety advantage could be attributed to NR-V04’s superior specificity as a PROTAC that targets NR4A1 rather than a collection of other known celastrol targets, which reduces off-target effects that could lead to toxicity in patients. One caveat related to the NR- V04 is the usage of VHL which is a well-established tumor suppressor gene^56^ and very commonly mutated in human cancers such as RCCs. We have been actively developing other E3-recruiting NR4A1 PROTACs, but at the current stage, none of those different PROTACs exhibit better tumor suppression and safety profile than NR-V04.

## Materials and Methods

### Chemistry

DMF and DCM were obtained via a solvent purification system by filtering through two columns packed with activated alumina and 4 Å molecular sieve, respectively. Water was purified with a Milli-Q Simplicity 185 Water Purification System (Merck Millipore). All other chemicals and solvents obtained from commercial suppliers were used without further purification. Flash chromatography was performed using silica gel (230–400 mesh) as the stationary phase.

Reaction progress was monitored by thin-layer chromatography (silica-coated glass plates) and visualized by 256 nm and 365 nm UV light, and/or by LC-MS. ^1^H NMR spectra were recorded in CDCl_3_ or CD_3_OD at 600 MHz, and ^13^C NMR spectra were recorded at 151 MHz using a Bruker (Billerica, MA) DRX Nuclear Magnetic Resonance (NMR) spectrometer. Chemical shifts δ are given in ppm using tetramethylsilane as an internal standard. Multiplicities of NMR signals are designated as singlet (s), doublet (d), doublet of doublets (dd), triplet (t), quartet (q), A triplet of doublets (td), A doublet of triplets (dt), and multiplet (m). All final compounds for biological testing were of ≥95.0% purity as analyzed by LC-MS, performed on an Advion AVANT LC system with the expression CMS using a Thermo Accucore™ Vanquish™ C18+ UHPLC Column (1.5 µm, 50 x 2.1 mm) at 40 °C. Gradient elution was used for UHPLC with a mobile phase of acetonitrile and water containing 0.1% formic acid. High resolution mass spectra (HRMS) were recorded on an Agilent 6230 Time-of-Flight (TOF) mass spectrometer.

Intermediate 1a to 3a were synthesized according to previously reported procedure^57^, briefly, to a solution of VHL-amine HCl salt (1.0 equiv), and corresponding acid-terminated linkers (1.0 equiv) in 5 mL DCM was added HATU (1.1 equiv) and DIPEA (5.0 equiv), the reaction was stirred at room temperature overnight. After extraction with EA and brine, the residue was concentrated and purified by silica gel column chromatography to yield 1a to 3a.

tert-butyl (2-(2-(2-(((S)-1-((2S,4R)-4-hydroxy-2-(((S)-1-(4-(4-methylthiazol-5- yl)phenyl)ethyl)carbamoyl)pyrrolidin-1-yl)-3,3-dimethyl-1-oxobutan-2-yl)amino)-2- oxoethoxy)ethoxy)ethyl)carbamate (1a) was obtained as colorless oil with the 93% yield: ^1^H NMR (600 MHz, Chloroform-*d*) δ 8.67 (s, 1H), 7.57 – 7.35 (m, 5H), 7.33 – 7.27 (m, 1H), 5.20 – 5.03 (m, 1H), 4.79 – 4.50 (m, 3H), 4.17 – 3.80 (m, 3H), 3.69 – 3.49 (m, 7H), 3.45 – 3.28 (m, 2H), 2.57 – 2.51 (m, 4H), 2.22 – 1.99 (m, 1H), 1.59 – 1.48 (m, 12H), 1.05 (s, 9H). ESI [M+H]^+^ = 690.4 tert-butyl ((S)-13-((2S,4R)-4-hydroxy-2-(((S)-1-(4-(4-methylthiazol-5- yl)phenyl)ethyl)carbamoyl)pyrrolidine-1-carbonyl)-14,14-dimethyl-11-oxo-3,6,9-trioxa-12-azapentadecyl)carbamate (2a) was obtained as colorless oil with the 91% yield: ^1^H NMR (600 MHz, Chloroform-*d*) δ 8.69 (s, 1H), 7.54 – 7.46 (m, 1H), 7.43 – 7.33 (m, 5H), 5.20 (s, 1H), 5.11 – 5.01 (m, 1H), 4.71 (t, *J* = 7.8 Hz, 1H), 4.63 – 4.53 (m, 1H), 4.55 – 4.46 (m, 1H), 4.08 – 3.93 (m, 3H), 3.73 – 3.61 (m, 9H), 3.54 (t, *J* = 5.4 Hz, 2H), 3.36 – 3.26 (m, 2H), 2.52 (s, 3H), 2.42 – 2.34 (m, 1H), 2.15 – 2.07 (m, 1H), 1.48 (d, *J* = 7.0 Hz, 3H), 1.43 (s, 9H), 1.06 (s, 9H). ESI [M+H]^+^ = 733.3 tert-butyl ((S)-16-((2S,4R)-4-hydroxy-2-(((S)-1-(4-(4-methylthiazol-5- yl)phenyl)ethyl)carbamoyl)pyrrolidine-1-carbonyl)-17,17-dimethyl-14-oxo-3,6,9,12-tetraoxa-15- azaoctadecyl)carbamate (3a) was obtained as colorless oil with the 91% yield: ^1^H NMR (600 MHz, Chloroform-*d*) δ 8.68 (s, 1H), 7.46 – 7.41 (m, 1H), 7.41 – 7.35 (m, 4H), 7.26 – 7.20 (m, 1H), 5.27 – 5.16 (m, 1H), 5.10 – 5.02 (m, 1H), 4.70 (t, *J* = 7.9 Hz, 1H), 4.59 (d, *J* = 8.7 Hz, 1H), 4.51 (s, 1H), 4.06 (s, 2H), 3.95 (d, *J* = 11.2 Hz, 1H), 3.80 (s, 1H), 3.71 – 3.59 (m, 12H), 3.56 (t, *J* = 5.2 Hz, 2H), 3.36 – 3.26 (m, 2H), 2.83 – 2.75 (m, 1H), 2.52 (s, 3H), 2.40 – 2.31 (m, 1H), 2.15 – 2.07 (m, 1H), 1.50 – 1.39 (m, 12H), 1.05 (s, 9H). ESI [M+H]^+^ = 778.4

General procedure for the synthesis of PROTACs: 1a, 2a, or -3a (1.0 equiv) was dissolved in 5 mL DCM/TFA (2:1) and stirred at room temperature for 6h. The resulting mixture was concentrated under vacuum and re-dissolved in DMF, celastrol was added followed by DIPEA (5.0 equiv) and HATU (1.1 equiv). The reaction mixture was then stirred at room temperature overnight. Water (20 mL) and EtOAc (40 mL) were added. The organic phase was washed with brine (10 mL), dried over Na_2_SO_4_, filtered, and concentrated. The residue was purified by silica gel column chromatography to yield NR-V02, NR-V03, and NR-V04, respectively.

NR-V02 was obtained as red solid (14 mg, 50% yield). ^1^H NMR (600 MHz, Chloroform-*d*) δ 8.67 (s, 1H), 7.40 – 7.33 (m, 6H), 7.08 (s, 1H), 7.03 (dd, *J* = 7.1, 1.4 Hz, 1H), 6.53 (s, 1H), 6.44 – 6.40 (m, 1H), 6.34 (d, *J* = 7.4 Hz, 1H), 5.11 – 5.07 (m, 1H), 4.72 (t, *J* = 8.1 Hz, 1H), 4.56 (d, *J* = 8.6 Hz, 1H), 4.52 (s, 1H), 4.12 – 3.96 (m, 3H), 3.76 – 3.13 (m, 12H), 2.52 (s, 3H), 2.47 – 2.39 (m, 2H), 2.20 (s, 3H), 2.17 – 1.76 (m, 7H), 1.69 – 1.41 (m, 11H), 1.24 (s, 3H), 1.16 – 0.97 (m, 16H), 0.61 (s, 3H). ^13^C NMR (151 MHz, CDCl_3_) δ 178.52, 178.35, 171.51, 171.10, 170.68, 169.89, 165.08, 150.32, 148.47, 146.11, 143.22, 134.68, 131.61, 130.84, 129.55, 127.28, 126.42, 119.48, 118.11, 117.71, 70.84, 70.14, 70.07, 58.69, 57.46, 56.77, 48.88, 47.40, 46.33, 45.11, 44.36, 43.14, 40.38, 39.39, 39.35, 38.19, 36.35, 35.94, 35.27, 34.87, 33.83, 33.50, 31.62, 31.05, 30.76, 29.96, 29.71, 29.38, 28.64, 26.53, 26.45, 26.39, 22.23, 21.68, 18.40, 16.11, 10.30, 8.72. MS m/z: [M+H]^+^ calculated for C_58_H_80_N_5_O_9_S_1_^+^: 1022.5671, found 1022.5680.

NR-V03 was obtained as red solid (12 mg, 51% yield). ^1^H NMR (600 MHz, Chloroform-*d*) δ 8.67 (s, 1H), 7.48 – 7.35 (m, 5H), 7.24 (d, *J* = 8.5 Hz, 1H), 7.11 – 7.05 (m, 1H), 7.02 (d, *J* = 7.0, 1.4 Hz, 1H), 6.53 – 6.50 (m, 1H), 6.37 – 6.30 (m, 2H), 5.14 – 5.05 (m, 1H), 4.72 (t, *J* = 7.9 Hz, 1H), 4.57 (d, *J* = 8.7 Hz, 1H), 4.52 (s, 1H), 4.09 – 4.00 (m, 3H), 3.87 – 2.98 (m, 16H), 2.53 (s, 3H), 2.50 – 2.39 (m, 2H), 2.20 (s, 3H), 2.15 – 1.81 (m, 7H), 1.69 – 1.36 (m, 11H), 1.25 (s, 3H), 1.12 (d, *J* = 16.9 Hz, 6H), 1.05 (s, 9H), 1.02 – 0.96 (m, 1H), 0.62 (s, 3H). ^13^C NMR (151 MHz, CDCl_3_) δ 178.46, 178.31, 171.44, 170.90, 170.53, 169.90, 165.02, 150.29, 148.48, 146.09, 143.31, 134.51, 131.63, 130.82, 129.54, 127.33, 126.45, 119.45, 118.10, 117.55, 70.67, 70.34, 70.31, 70.12, 69.87, 67.99, 58.60, 57.33, 56.72, 48.84, 47.40, 45.11, 44.37, 43.09, 40.29, 39.34, 39.29, 38.25, 36.35, 35.89, 35.20, 34.92, 33.79, 33.54, 31.62, 31.09, 30.78, 29.99, 29.71, 29.36, 28.68, 26.51, 26.40, 25.62, 22.23, 21.70, 18.38, 16.12, 10.29, 8.73. MS (ESI); m/z: [M+H]^+^ calculated for C_60_H_84_N_5_O_1_S_1_^+^: 1066.5933, found 1066.5933.

NR-V04 was obtained as red solid (24 mg, 55% yield). ^1^H NMR (600 MHz, Chloroform-*d*) δ 8.67 (s, 1H), 7.43 – 7.35 (m, 5H), 7.17 (s, 1H), 7.04 (d, *J* = 7.0 Hz, 1H), 6.53 (s, 1H), 6.45 (s, 1H), 6.34 (d, *J* = 7.2 Hz, 1H), 5.12 – 5.05 (m, 1H), 4.68 (t, *J* = 8.0 Hz, 1H), 4.57 (d, *J* = 8.9 Hz, 1H), 4.51 (s, 1H), 4.09 – 3.92 (m, 3H), 3.73 – 3.15 (m, 17H), 2.53 (s, 3H), 2.48 – 2.38 (m, 2H), 2.21 (s, 3H), 2.16 – 1.80 (m, 12H), 1.69 – 1.42 (m, 13H), 1.25 (s, 3H), 1.12 (d, *J* = 14.7 Hz, 6H), 1.06 – 0.92 (m, 10H). ^13^C NMR (151 MHz, CDCl_3_) δ 179.35, 178.35, 171.61, 171.14, 170.37, 170.25, 165.37, 150.30, 148.41, 146.17, 143.34, 135.18, 131.68, 130.73, 129.52, 127.21, 126.53, 119.36, 118.24, 118.06, 70.89, 70.04, 69.89, 69.66, 69.51, 59.04, 57.30, 56.84, 48.89, 45.20, 44.29, 43.20, 40.44, 39.31, 39.12, 38.19, 36.31, 36.23, 35.51, 34.65, 33.83, 33.55, 31.57, 30.79, 29.75, 29.27, 28.65, 26.36, 22.09, 21.61, 18.30, 16.11, 10.30. MS (ESI); m/z: [M+H]^+^ calculated for C_62_H_88_N_5_O_11_S_1_^+^: 1110.6196, found 1110.6199.

### Single-cell RNA analysis

We utilized multiple datasets from GEO (GSE12057552, GSE14819053, GSE15880354, GSE12163850) and the human TCGA melanoma data cohort for analysis. Data visualization and analysis were conducted using Single Cell Portal, Seurat R package (V2.3.4), and Gene Set Enrichment Analysis (GSEA). The code used in our analysis is available on GitHub (https://github.com/Levy0803/NR4A1/projects?query=is%3Aopen).

### Cell lines and cell culture

Human melanoma cell lines: A375 and CHL-1 (gift from Dr. Zhou Daohong’s lab) were cultured in Dulbecco’s Modified Eagles’ Medium (DMEM, D5796, Sigma), WM164 and M229 (gift from Dr. Smalley Keiran’s lab) were cultured in RPMI-1640 (R8758, Sigma), B16F10 (CRL-6475, ATCC). Mouse melanoma cell lines: SW1 and SM1 (gift from Dr. Smalley Keiran’s lab) were cultured in RPMI. Human epithelial kidney cell lines: HEK293T (293T, CRL-3216, ATCC) and Vhl-KO-293T(gift from Dr. Zhou Daohong’s lab) were cultured in DMEM. Mouse colon cancer cell lines: MC38 (ENH204-FP, Kerafast Inc.) was cultured in DMEM supplemented with 1mM glutamine, 0.1 M non-essential amino acids (11140050, Thermo Fisher), 1 mM sodium pyruvate (11360070, Thermo Fisher), 10 mM HEPES (15630080, Thermo Fisher). All cell culturing medium were supplemented with 10% Fetal Bovine Serum (FBS, F2442, Sigma) and 100LU/ml penicillin, and 100 µg/ml streptomycin (P4333, Sigma).

### Immunoblotting

Samples were lysed using RIPA lysis buffer (150 mM NaCl, 5 mM EDTA, 50 mM Tris pH 8.0, 1% sodium deoxycholate, 1% NP-40, 0.5% SDS) supplemented with 1 mM dithiothreitol and protease inhibitors. The lysates were separated by SDS-PAGE and analyzed using standard western blotting procedures^58^. Antibodies: human NR4A1 (ab153914, Abcam, 1:1000), mouse NR4A1 (14-5965-82, Invitrogen, 1:1000), NR4A2 (AV38753, Sigma, 1:1000), NR4A3 (TA804893, Thermo Fisher, 1: 1000), VHL (68547,Cell Signaling, 1:1000), β-actin (13E5, Cell Signaling, 1:5,000), β-tubulin (9F3, Cell Signaling, 1: 5000), GAPDH (D16H11, Cell Signaling, 1: 5000) were employed for protein detection.

### Cell transfection

HEK293T cells were transfected with 3 µg Flag-NR4A1 (HG17699-CF, SinoBiological, sequencing primer is T7(TAATACGACTCACTATAGGG) BGH(TAGAAGGCACAGTCGAGG), Flag-tag primer is GATTACAAGGATGACGACGATAAG) for 36 hours in Opti-MEM (31985062, Thermo Fisher) medium with 10 µL GeneTran III reagent (GT2211, Biomiga) in 6 cm^2^ cell culture dishes.

### Co-immunoprecipitation

HEK293T cells were transfected with Flag-NR4A1 for 36 hours. Afterward, the cells were treated with either DMSO or 500 μM NR4A1 for 16 hours, and 500 nM MG132 was included to prevent protein degradation. Proteins were extracted using RIPA lysis buffer without SDS (150 mM NaCl, 5 mM EDTA, 50 mM Tris pH 8.0, 1% sodium deoxycholate, 1% NP-40) supplemented with 1% protease inhibitor cocktail (C0001, TargetMol). Immunoprecipitation was performed using Anti-FLAG M2 Magnetic Beads (M8823, Sigma-Aldrich) as per the manufacturer’s protocol. The immunoprecipitated samples were eluted with 2× SDS sample buffer and boiled for 5 minutes at 95°C. The samples were then subjected to denaturation, SDS–polyacrylamide gel electrophoresis (SDS–PAGE), and immunoblotting analysis.

### Proximity ligation assay (PLA)

5 × 10^4^ CHL-1 and A375 cells were cultured in Nunc™ Lab-Tek™ Chamber Slide (177402, Thermos Fisher). The cells were treated with DMSO, 500 nM celastrol, and 500 nM NR-V04 for 16 hours with MG132 inhibitor. The proximity ligation assay (PLA) was conducted with Duolink^®^ In Situ Red Starter Kit Mouse/Rabbit (DUO92101, Sigma) following the standard protocol. Additionally, a non-stain cell group was included as a negative control to account for background signal during PLA assay. The signal was detected using a Zeiss LSM 710 confocal microscope. Primary antibody: NR4A1 (HPA059742, Sigma, 2 μg/mL, isotype-rabbit), VHL (MA5-13940, Thermo Fisher, 2 μg/mL, isotype-mouse).

### Quantitative PCR (qPCR)

Total RNA was extracted from cells using RNeasy Kits (74004, QIAGEN) following the manufacturer’s instructions. cDNA synthesis was performed using SuperScript™ III Reverse Transcriptase (18080044, Thermo Fisher) with 100 ng/μL RNA as the template. Amplification of cDNA was carried out using 2X PowerUP SYBR Green Master Mix (Thermo Fisher, A25778). Each reaction was conducted in triplicates, and results were normalized to the expression of a housekeeping gene. Primer (Eurofins) : human β*-actin* (Forward Sequence: CACCATTGGCAATGAGCGGTTC; Reverse Sequence: AGGTCTTTGCGGATGTCCACGT), human *NR4A1* (Forward Sequence: GGACAACGCTTCATGCCAGCAT; Reverse Sequence: CCTTGTTAGCCAGGCAGATGTAC).

### Pharmacokinetic study of NR-V04

The pharmacokinetic study of NR-V04 subjected to PK studies was done by Bioduro Inc. on healthy male C57BL/6 mice weighing between 20 g and 25 g with three animals in each group. Test compound (*i.v.*) was dissolved in 5% DMSO/3% tween 80 in PBS with 0.5 mEq 1N HCl and was administered (n = 3 per group) at a dose level of 2 mg/kg (concentration: 0.4 mg/mL). Test compounds (*i.p.*) were dissolved in 5% DMSO/3% tween 80 in PBS with 2 mEq 1N HCl (concentration 4 mg/kg) and were administrated at a dose level of 20 mg/kg. Plasma samples (*i.v.*) were collected from the orbital plexus at 0.083, 0.25, 0.5, 1.0, 2.0, 4.0, 8.0 and 24 h, and plasma samples (*i.p.*) were collected from the orbital plexus at 0.25, 0.50, 1.0, 2.0, 4.0, 8.0, and 24 h postdose. The drug concentrations in the samples were quantified by liquid chromatography/tandem mass spectrometry (LC-MS/MS). Pharmacokinetic parameters were calculated from the mean plasma concentration by non-compartmental analysis.

### B cell proliferation assay

B cells were isolated from the spleen by using EasySep™ Mouse B Cell Isolation Kit (19854, STEMCELL) and cultured in RPMI supplemented with 0.1 M non-essential amino acids (11140050, Thermo Fisher), 1 mM sodium pyruvate (11360070, Thermo Fisher), 1mM Glutamax, 10 mM HEPES (15630080, Thermo Fisher), 0.1% 2-mercaptoethanol (21985023, Thermo Fisher), 10% Fetal Bovine Serum (FBS, F2442, Sigma) and 100LU/mL penicillin, and 100 µg/mL streptomycin (P4333, Sigma). B cells were stained with 5 nM CellTrace™ Violet Cell (C34557, Thermo Fisher) for 8 minutes in 37 °C water bath. B cells were plated at a concentration of 5L×L10^5^ cells per 200 μL in round-bottom 96-well plates treated with 0.1% B16F10 melanoma cell lysis or 10 µg/mL goat anti-mouse IgM F(ab’)2 (115-006-020, Jackson ImmunoResearch). B cells were treated with NR-V04 or DMSO for 24 hours and collected for flow cytometry analysis.

### Animal Experiment

All animal studies were approved by the University of Florida Institutional Animal Care and Use Committee (IACUC) under protocol 202110399. All animals were housed in a pathogen-free Association for Assessment and Accreditation of Laboratory Animal Care accredited facility at the University of Florida and performed in accordance with IACUC guidelines. The room temperature is 21.1–23.3 °C with an acceptable daily fluctuation of 2 °C. Typically the room is 22.2 °C all the time. The humidity set point is 50% but can vary ±15% daily depending on the weather. The photoperiod is 12:12 and the light intensity range is 15.6–25.8 FC.

7 to 9-week-old C57BL/6J, B6.129S2-*Ighm*^tm1Cgn^/J (002288), B6.129S2-*Nr4a1^tm1Jmi^*/J (NR4A1^-/-^, 006187), NOD-SCID interleukin-2 receptor gamma null (NSG) mice were purchased from Jackson Laboratories (Bar Harbor, ME). For all studies involving NR-V04 and celastrol, the compounds were formulated in 50% phosal PG, 45% miglyol 810N, and 5% polysorbate 80 and administered via IP injection.

For syngeneic tumor models, 5L×L10^5^ cells MC38, Yumm 1.7 and Yummer 1.7, 1L×L10^5^ cells of B16F10, were resuspended in phosphate buffered saline (PBS), and implanted into 7 to 9- week-old mice subcutaneously. For tumor inhibition experiment, mice were treated with 1.8 mg/kg NR-V04 or 0.75 mg/kg celastrol by IP injection twice weekly. For immune profile experiment, once tumors reached 0.5 cm in diameter, mice were treated with 1.8 mg/kg NR- V04 by IP injection twice weekly. For toxicity experiment, male and female non-tumor bearing mice were treated with 2 mg/kg NR-V04 for two doses for first week and 5 mg/kg for another two doses in second week. Blood samples were taken from the submandibular (facial) vein before each treatment and 42 days later for hematological analysis and body weight were measured twice per week up to day 42. Mice were euthanized in accordance with IACUC protocol once the largest tumors reached 2 cm in diameter and tissues were collected for further analysis.

### Flow cytometry

Mouse tumors were excised and ∼200 mg of tumor tissue was enzymatically and mechanically digested using the mouse Tumor Dissociation Kit (Miltenyi Biotec) to obtain a single-cell suspension. Human tumor samples and sections were enzymatically and mechanically digested using the human Tumor Dissociation Kit (Miltenyi Biotec) to obtain single-cell suspension. Red blood cells were lysed using ACK lysis buffer and mononuclear cells were isolated by density gradient using SepMate Tubes (StemCell Technologies) and Lymphoprep density gradient media (StemCell Technologies). Mouse cells were then washed and incubated with combinations of the following antibodies: anti-mouse CD62L-BV785 (clone MEL-14), anti-mouse MHCII I-A/I-E-BB515 (Clone 2G9, BD Biosciences, 1:400), anti-mouse CD11B-PEdazzle 594 (clone M1/70, 1:200), anti-mouse CD45-AF532 (clone 30F.11), anti-mouse CD3-APC/Cy7(clone 17A2), anti-mouse CD8-BV510 (clone 53-6.7), anti-mouse CD4-BV605 (clone GK1.5), anti-mouse NK1.1-AF700 (clone PK136), anti-mouse/human CD45R/B220-BV 570 (clone RA3-6B2), anti-mouse CD138-BV711 (clone 281-2), anti-mouse IgD-BV711 (clone 281-2), anti-mouse IgD- PE-Cy7 (clone 11-26c.2a), anti-mouse Ly6G-FITC (clone IA8), anti-mouse Ly6C-BV711 (clone HK1.4), anti-mouse CD38-PerCP/Cy 5.5 (clone 90) anti-mouse IgM-AF488 (clone RMM-1), anti-mouse CD25-PE-Cy5 (clone PC61) plus zombie red (cell viability, Thermo Fisher) and mouse FcR blocker (anti-mouse CD16/CD32, clone 2.4G2, BD Biosciences). After surface staining, cells were fixed and permeabilized using the FOXP3/Transcription Factor Staining Buffer Set (eBioscience). Cells were stained with a combination of the following antibodies: anti-mouse FOXP3-efluor 450 (clone FJK-16S, 1:50, eBioscience), anti-mouse Granzyme B-BV421 (clone GB11), anti-mouse NR4A1-PE (clone 12.14, eBioscience). Human cells were stained with a combination of the following antibodies: anti-human CD45-BV510 (clone H130), anti-human CD3-AF700 (clone HIT5a), anti-human CD4-BV421 (clone OKT4), anti-human CD8-BV711 (clone RPA-T8), anti-human CD127-BV605 (clone A019DS), anti-human CD25-PE-Cy7 (clone MA251) plus FVD-eFluor-780 (eBioscience) and human FcR blocking Reagent (StemCell Technologies). Cells were washed then fixed and permeabilized using the eBioscience FOXP3/Transcription Factor Staining Buffer Set. Cells were further stained with a combination of the following antibodies: anti-human FOXP3-FITC (clone 206D), anti-human NR4A1-PE (clone D63C5, Cell Signaling). Flow cytometry was performed on a 3 laser Cytek Aurora Cytometer (Cytek Biosciences, Fremont, CA) and analyzed using FlowJo software (BD Biosciences). All antibodies are from Biolegend, unless otherwise specified. Most antibodies were used at 1:100 dilution for flow cytometry, unless otherwise specified.

### Hematology analysis

Mouse blood was prepared in micro-centrifuge tubes containing PBS with 10 mM EDTA. Blood indices were analyzed using an automated hematology analyzer (Element HT5, Heska).

### H&E staining

H&E staining for mouse tissues were fixed in 10% formalin (SF98-4, Thermo Fisher) processed in UF pathology core. H&E staining was conducted using standard procedures. The sections were hydrated through xylene and a series of ethanol, followed by staining with Hematoxylin and Eosin on slides.

### Statistics

Graphs and statistical analyses were performed using Prism software (GraphPad Software) unless otherwise specified. Tumor growth curves were compared using a two-way analysis of variance (ANOVA). For comparisons involving three or more groups, a one-way ANOVA was conducted, followed by Dunnett’s multiple comparison test for specific group comparisons.

Unpaired T tests were used to compare means between two groups.

## Author Contributions

Funding: W.Z. and G.Z.; Conception and design: W.Z., G.Z., L.W., Y.X.; Development of methodology: L.W., Y.X., Y.L., R.P.M., J.M., M-C.K., Y.L., E.M.; Acquisition of cancer specimens and data: L.W., Y.X., Y.L., R.P.M., J.M., M-C.K., Y.L., U.M.P., W.Z.; Analysis and interpretation of data: L.W., Y.X., X.L., D.S., K.G., G.Z. W.Z.; Writing, review, and/or revision of the manuscript: L.W. and Y.X. for first draft; W.Z., G.Z., K.S.S., D.Z. for rewriting, revision and editing. Supervision: W.Z., G.Z.

## Acknowledgements

The work is mainly supported by DOD/CDMRP grant BC200100 (Partnering PIs: W.Z. and G.Z.) and an iAward from Sanofi (PI: W.Z.). The work is partially supported by NIH grants CA200673 (W.Z.), CA203834 (W.Z.), CA260239 (W.Z.), and DOD/CDMRP grant BC200100 (W.Z.). W.Z. was also supported by an endowment fund from the Dr. and Mrs. James Robert Spenser Family.

## Supplementary Information

**Figure S1.**
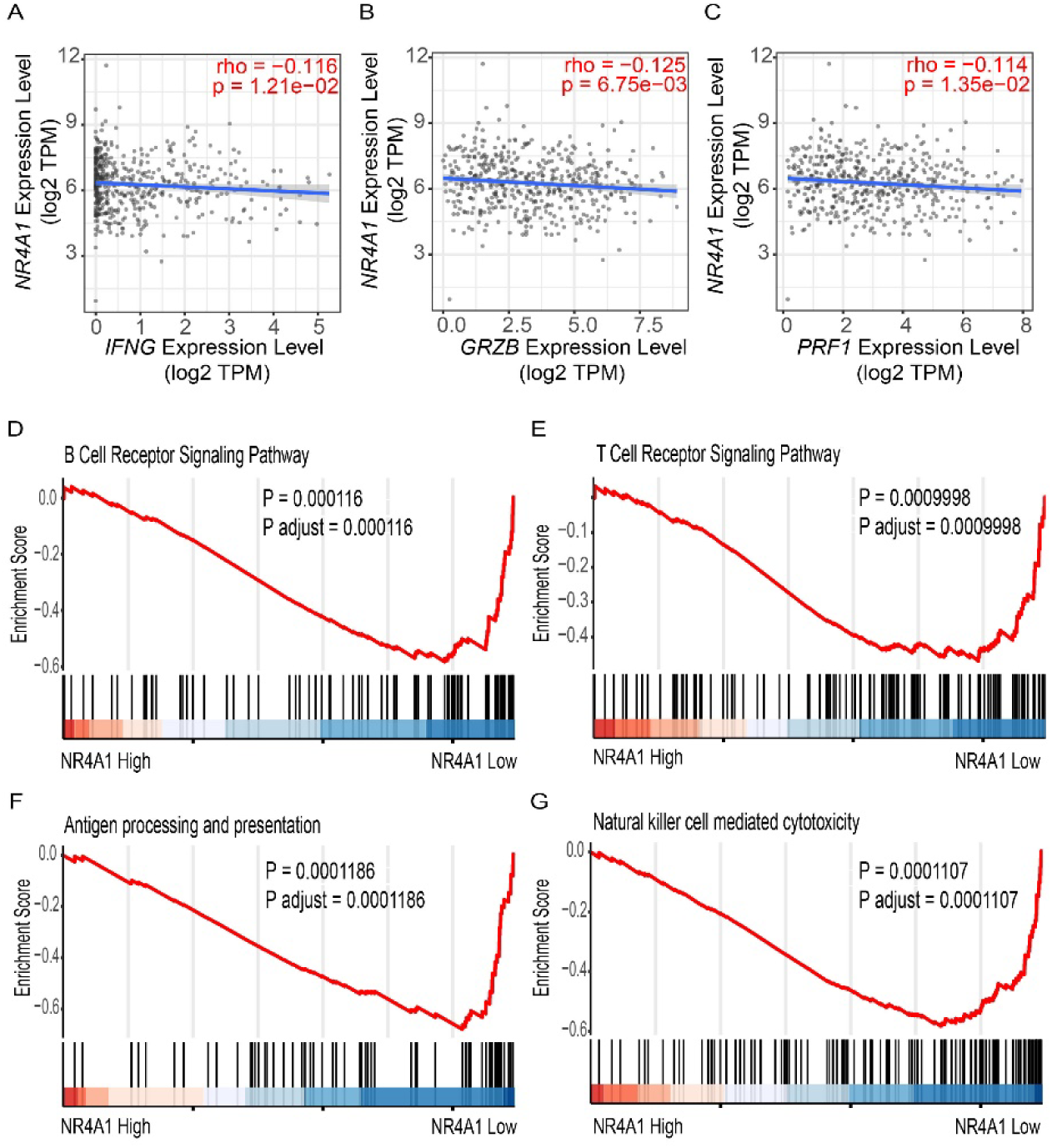
NR4A1 expression is inversely correlated with effector molecules required for T cell activation. **A-C.** NR4A1 expression is inversely correlated with (A) IFNG, (B) GZMB, and (C) PRF1 gene expression in the TCGA melanoma datasets. **D-G.** GSEA analysis showing the enrichment of immune activation pathways in the melanoma specimens with low NR4A1 expression. The TCGA human melanoma dataset was divided into tertiles based on NR4A1 expression. GSEA analysis was performed using the highest tertile versus the lowest tertile of NR4A1 expression. Supplementary to Figure 1.

**Figure S2.**
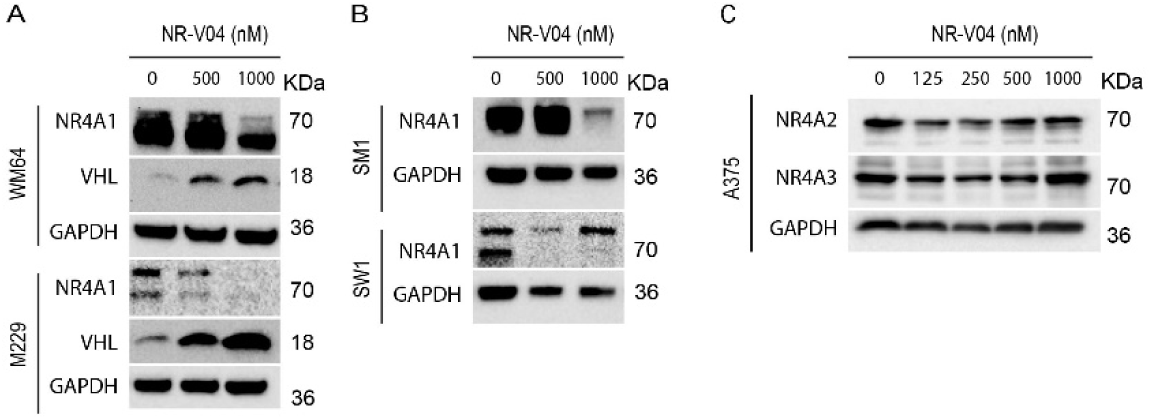
NR-V04 induces NR4A1 degradation. **A**. NR-V04 effectively promoted the degradation of NR4A1 in two more human melanoma cell lines in 16 hours, including WM164 and M229, while simultaneously stabilizing VHL expression. **B**. NR-V04 effectively promoted the degradation of NR4A1 in two mouse melanoma cell lines in 16 hours, including SM1 and SW1. **C.** NR-V04 did not induce the degradation of NR4A2 and NR4A3. n=2 for A-C. Supplementary to Figure 4.

**Figure S3.**
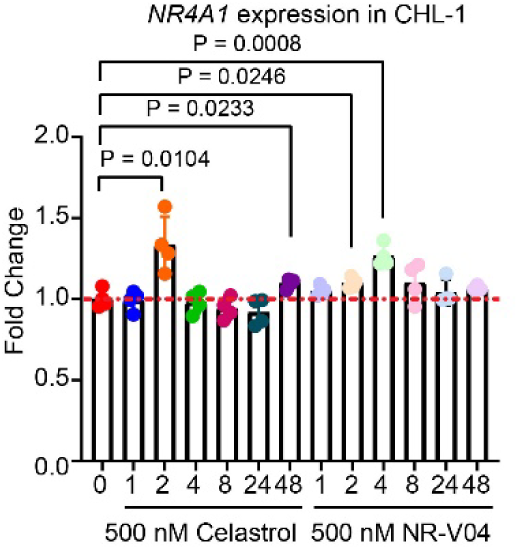
NR-V04 induces a transient elevation of *NR4A1* mRNA. Time-dependent degradation of *NR4A1*. CHL-1 cells were treated with 500nM of celastrol or NR-V04 at the indicated time points. RNA was prepared for reverse transcription and qPCR. *ACTB* was used as control. n = 3. Supplementary to Figure 4.

**Figure S4.**
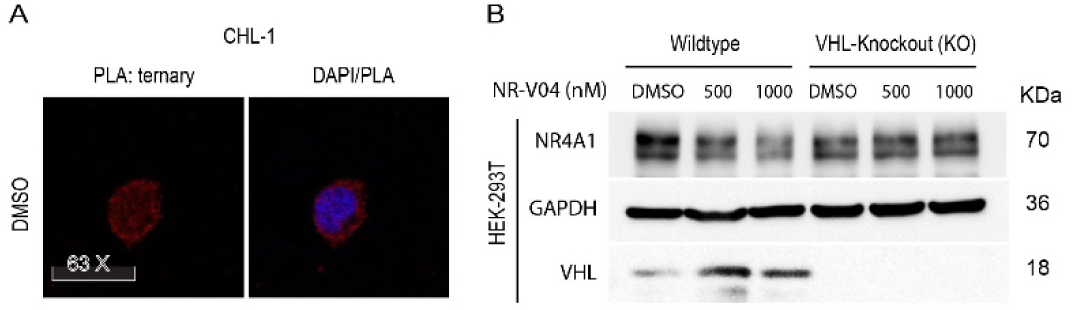
NR-V04 leads to a ternary formation and mediates NR4A1 degradation through the ubiquitin-proteasome system (UPS). **A**. Ternary complex formation was observed in CHL-1 cells with 16-hour treatments of 500 nM NR-V04, rather than DMSO or 500 nM celastrol, as detected by PLA (60 X magnification). **B.** NR4A1 degradation by 500 or 1000 nM of NR-V04 for 16 hours in the WT or VHL knockout HEK293T cells. Supplementary to Figure 5.

**Figure S5.**
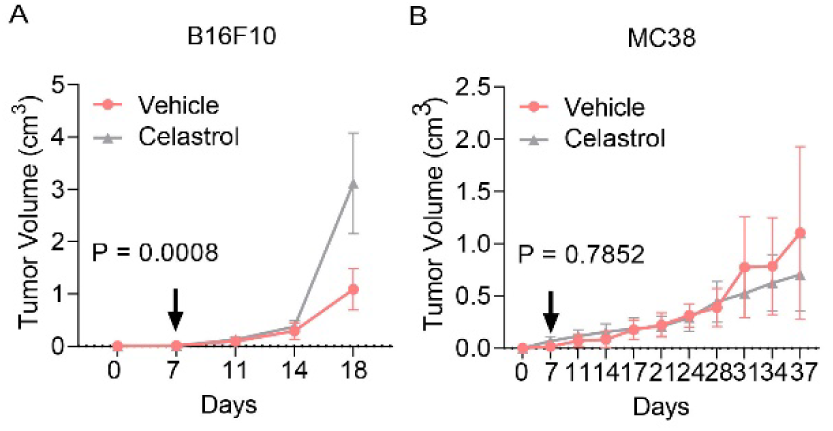
Celastrol did not inhibit tumor growth at equivalent doses. **A-B.** B16F10 (A) or MC38 (B) tumor-bearing mice were treated with vehicle or celastrol using the same treatment regimen as NR-V04 (Fig. 7). Supplementary to Figure 7.

**Figure S6.**
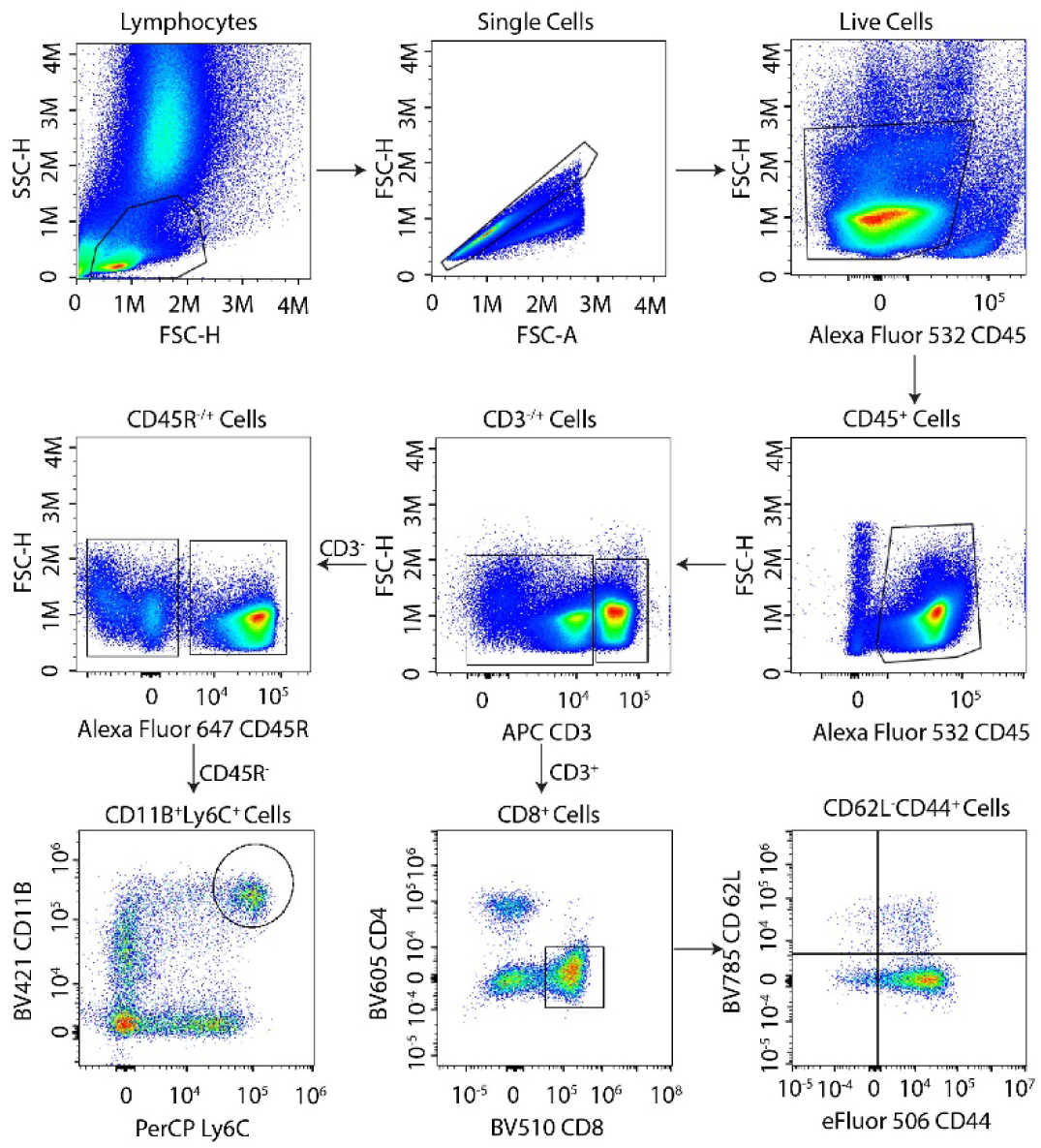
Schematic showing gating strategy for flow cytometry. Supplementary to Figure 8.

**Figure S7.**
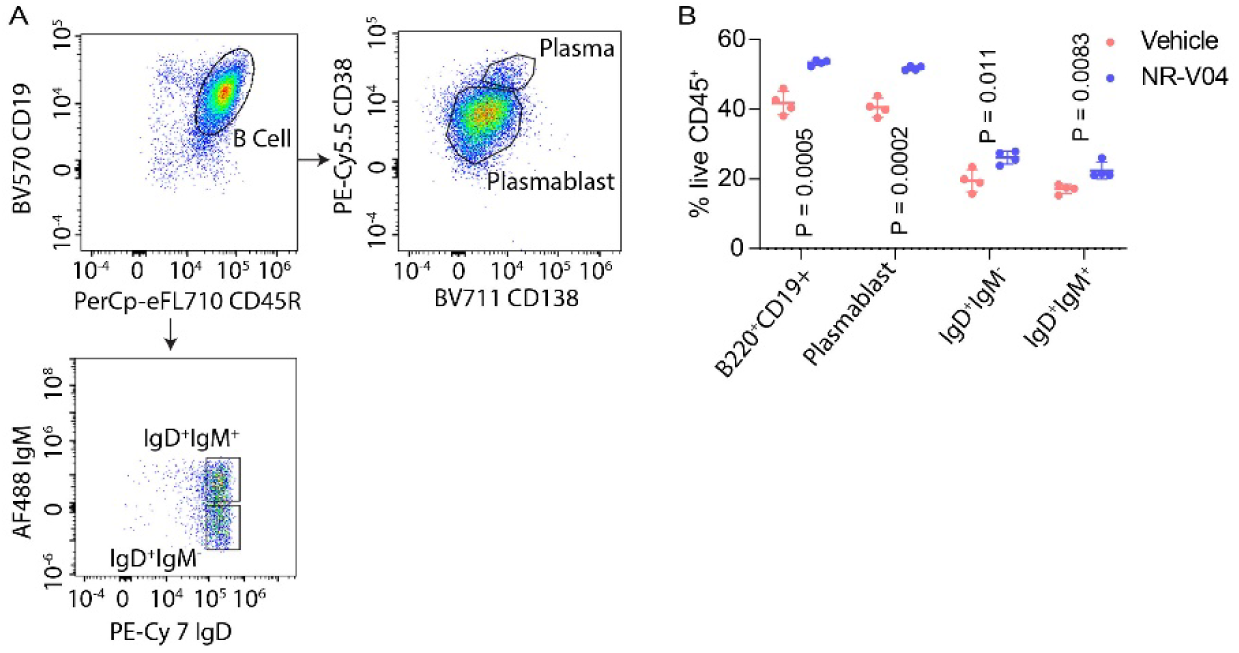
NR-V04 induces TI-B cells. **A.** Schematic showing gating strategy for flow cytometry of B cell populations. **B.** NR-V04 induces significant B cell proliferation and activation in B16F10 tumors. B16F10 tumor-bearing mice were treated with vehicle and NR-V04 as in Fig. 8. n = 7. Supplementary to Figure 8.

**Figure S8.**
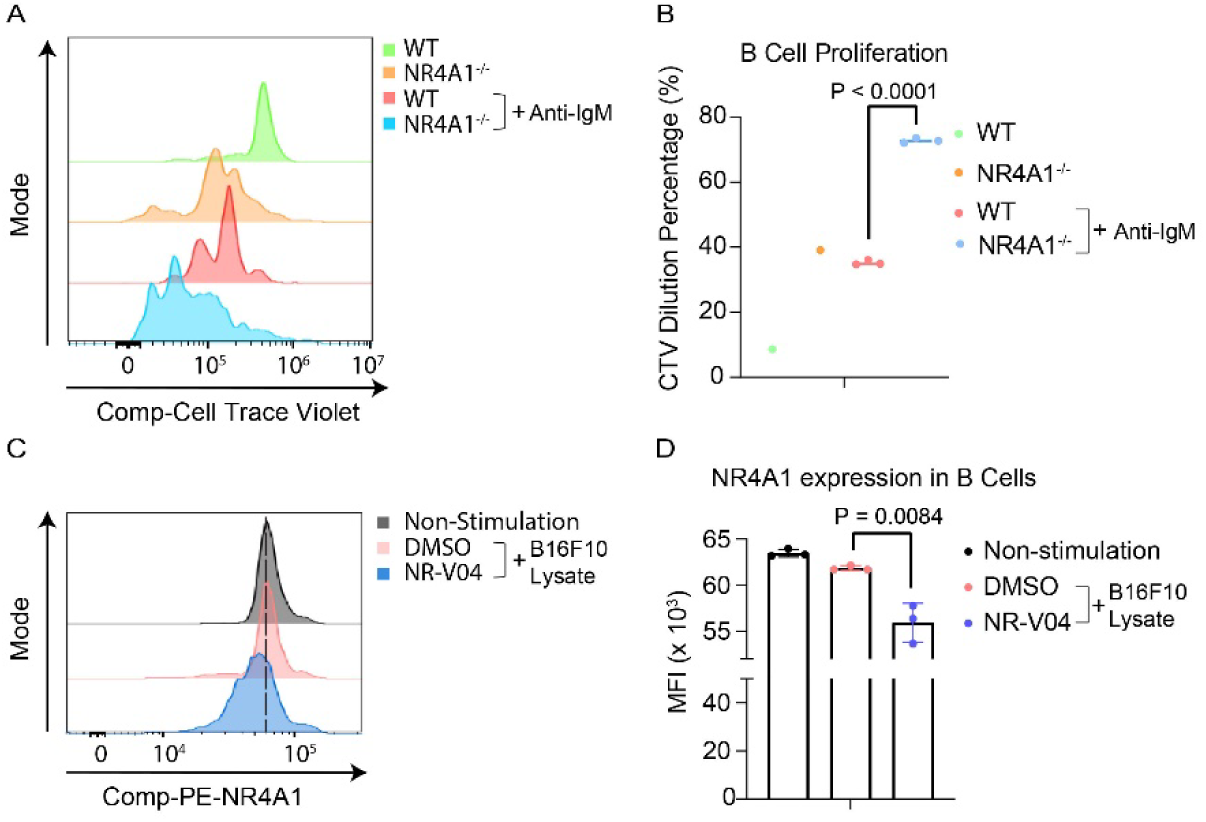
NR4A1 is critical in limiting B cell expansion. **A-B.** Splenic B cells from WT or *NR4A1*^-/-^ mice were either untreated or treated with IgM to induce B cell proliferation that was detected using flow cytometry of dilution of Cell Trace Violet. n = 3. **C-D.** Splenic B cells were treated with or without B16F10 lysis stimulation in vitro and B cell proliferation was assayed similarly using flow cytometry, with or without the presence of 500 nM NR-V04 treatment for 24 hours (n=3). (C) NR4A1 protein was determined by flow cytometry and (D) statistical summary. Supplementary to Figure 8.

**Figure S9.**
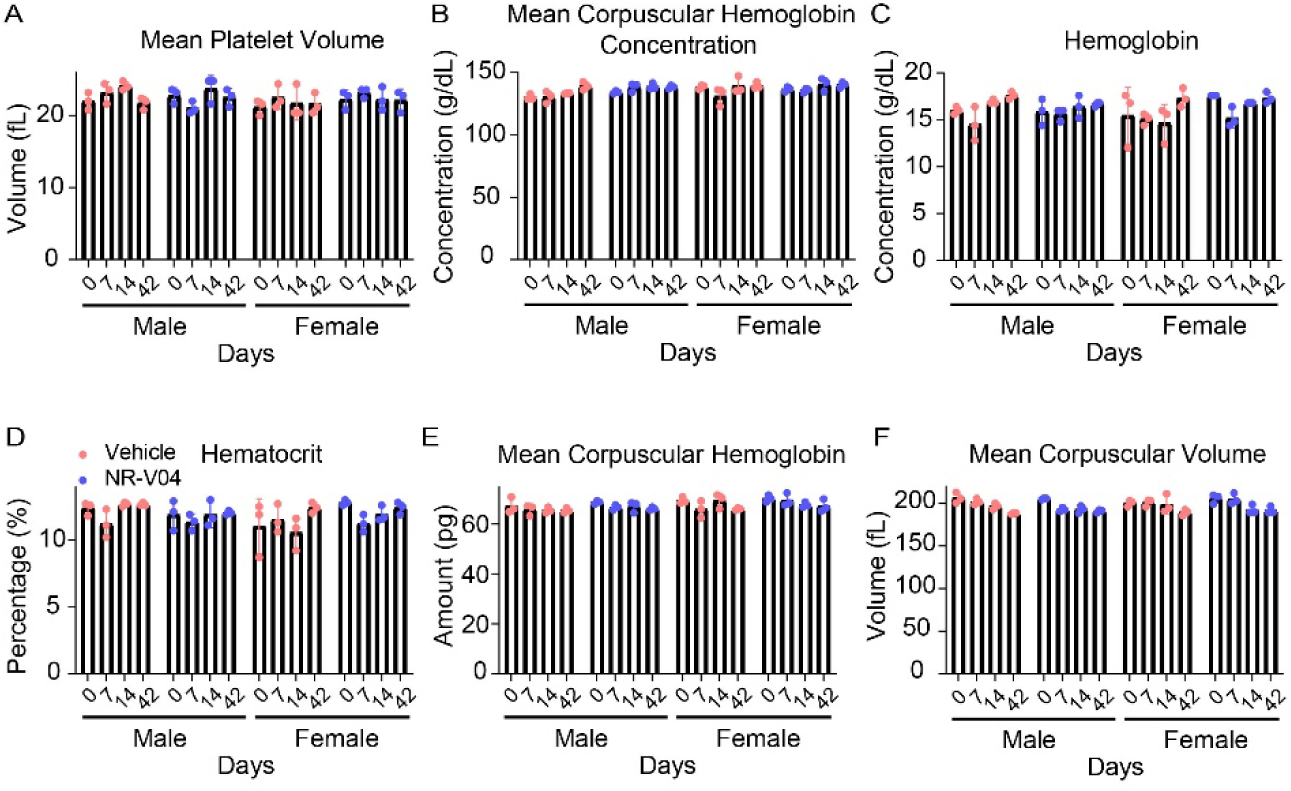
NR-V04 has minimal toxicity. **A-F.** Hematology analysis of different blood cell components after NR-V04 or vehicle treatment. NR-V04 did not cause significant changes in the hematologic profile including (A) mean platelet volume, (B) hemoglobin, (C) mean corpuscular hemoglobin, (D) mean corpuscular hemoglobin concentration, (E) hematocrit, and (F) mean corpuscular volume. n = 3 mice per group. Supplementary to Figure 9.

## Notes

### Competing Interest Statement

This project was partially supported by an iAward from Sanofi. W.Z., G.Z., D.Z., Y.X., and L.W. hold a patent related to the compound used in this study. X.L., D.S., K.R.G. are Sanofi Employees and holding Sanofi stocks.

